# Genetic load and adaptive potential of a recovered avian species that narrowly avoided extinction

**DOI:** 10.1101/2022.12.20.521169

**Authors:** Georgette Femerling, Cock van Oosterhout, Shaohong Feng, Rachel M. Bristol, Guojie Zhang, Jim Groombridge, M. Thomas P. Gilbert, Hernán E. Morales

**Affiliations:** Globe Institute, Faculty of Health and Medical Sciences, University of Copenhagen, Copenhagen, Denmark; Centro de Ciencias Genómicas, Universidad Nacional Autónoma de México, Cuernavaca, México; School of Environmental Sciences, University of East Anglia, Norwich, UK; Center for Evolutionary & Organismal Biology, Zhejiang University School of Medicine, Hangzhou, China; Liangzhu Laboratory, Zhejiang University Medical Center, 1369 West Wenyi Road, Hangzhou, China; Innovation Center of Yangtze River Delta, Zhejiang University, Jiashan, China; La Batie, Beau Vallon, Mahe, Seychelles; Durrell Institute of Conservation and Ecology, School of Anthropology and Conservation, Division of Human and Social Sciences, University of Kent, Canterbury, Kent, CT2 7NR, United Kingdom; University Museum, NTNU, Trondheim, Norway

## Abstract

High genetic diversity is often a good predictor of long-term population viability, yet some species persevere despite having low genetic diversity. Here we study the genomic erosion of the Seychelles paradise flycatcher (*Terpsiphone corvina*), a species that narrowly avoided extinction after having declined to 28 individuals in the 1960s. The species recovered unassisted to over 250 individuals in the 1990s and was downlisted from Critically Endangered to Vulnerable in the IUCN Red List in 2020. By comparing historical, pre-bottleneck (130+ years old) and modern genomes, we uncovered a 10-fold loss of genetic diversity. The genome shows signs of historical inbreeding during the bottleneck in the 1960s, but low levels of recent inbreeding after the demographic recovery. We show that the proportion of severely deleterious mutations has reduced in modern individuals, but mildly deleterious mutations have remained unchanged. Computer simulations suggest that the Seychelles paradise flycatcher avoided extinction and recovered due to its long-term small N_e_. This reduced the masked load and made the species more resilient to inbreeding. However, we also show that the chronically small N_e_ and the severe bottleneck resulted in very low genetic diversity in the modern population. Our simulations show this is likely to reduce the species’ adaptive potential when faced with environmental change, thereby compromising its long-term population viability. In light of rapid global rates of population decline, our work highlights the importance of considering genomic erosion and computer modelling in conservation assessments

## Introduction

Global population abundance of 4,392 species monitored over the last 40 decades has declined by 68% (Almond et al. 2022), threatening their long-term viability. On the IUCN Red List, 33,777 species (47.4%) are facing population decline, compared to 36,264 (50.9%) with stable population size, and 1274 (1.8%) that are increasing in size (IUCN 2022). A growing number of species are being classified as threatened with extinction, i.e., in the Red List categories of Vulnerable, Endangered, or Critically Endangered (Monroe et al. 2019). On the other hand, effective conservation management has been able to recover the population size after a severe bottleneck for a small number of species, resulting in their downlisting on the IUCN Red List (e.g., the snow leopard, the giant panda and the pink pigeon; (Mallon and Jackson 2017; Swaisgood et al. 2018). However, even when effective conservation actions are capable of reverting population declines, the negative genetic effects that may arise during population declines can be persistent (Tilman et al. 1994; Kuussaari et al. 2009). Populations that have recovered from a bottleneck could be subjected to a genetic drift debt in which they continue to lose genetic diversity, even after demographic recovery (Gilroy et al. 2017; Pinto et al. 2022). Population decline generates genetic drift and inbreeding that erode genetic diversity, compromising the viability of wild populations (Lynch et al. 1995; Lande and Shannon 1996; Willi et al. 2006; Bozzuto et al. 2019). Thus, investigating the evolutionary genomic consequence of population decline in species that have collapsed, but recovered and avoided extinction, is instrumental for better understanding and predicting extinction risk and the long-term viability of wild populations.

Empirical and simulation studies have shown that population bottlenecks and long-term small effective population sizes (N_e_) could be conducive to the reduction of deleterious variation through the purging of recessive, highly deleterious mutations (Dussex et al. 2021; Grossen et al. 2020; Hedrick and Garcia-Dorado 2016; Khan et al. 2021; Kleinman-Ruiz et al. 2022; van Oosterhout et al. 2022). Theoretically, this could make species more robust to inbreeding depression. However, small population sizes may also lead to the accumulation of genetic load through the accumulation of mildly deleterious mutations (Bertorelle et al. 2022; Grossen et al. 2020; Smeds and Ellegren 2022). Furthermore, genetic drift in small populations leads to reduced adaptive potential in the face of environmental change (Willi et al. 2006). At present, we have an incomplete understanding of the short- and long-term consequences of population decline and small effective population size on the viability and extinction risk of species (Hedrick and Garcia-Dorado 2016; Mable 2019; Forester et al. 2022).

The rate of genomic erosion and its impact on extinction probability is a complex outcome of the interaction between long-term trends of N_e_, recent population decline, the response of different types of genetic variation (e.g., deleterious mutations and adaptive genetic variation), and the rate of environmental change. Here, we quantify the genomic erosion in the Seychelles paradise flycatcher (*Terpsiphone corvina*), a species whose population declined to 28 individuals in 1965, followed by an (unassisted) recovery to over 250 individuals by the year 2000. In 2008 a self-sustaining, growing population was established on Denis Island with translocated individuals. After these demographic gains, the species’ conservation status in the IUCN Red List was downlisted from Critically Endangered to Vulnerable (IUCN 2022/1). We directly compare genomic variation pre- and post-population decline by sequencing whole genomes of museum-preserved samples (>130 years old) and modern samples. We show that the species suffered a 10-fold decline in genome-wide genetic diversity, one of the largest losses compared to other birds with reported historical comparisons. This has left the modern Seychelles paradise flycatcher population with a lower genome-wide diversity compared to other Endangered and Critically Endangered bird species. We used individual-based genomic simulations to investigate how the Seychelles paradise flycatcher managed to avoid extinction after suffering such a drastic population decline and loss of genetic diversity. Our results indicate that the ancestral, pre-bottleneck population had a reduced masked load due to their long-term small N_e_. This was conducive to less inbreeding depression that allowed them to avoid extinction and successfully recover. However, we also show that this long-term small N_e_, together with the substantial genetic diversity loss, have likely reduced the species’ adaptive potential and jeopardised their long-term viability when faced with environmental change.

## Methods

### Study system and sampling

The Seychelles paradise flycatcher historically inhabited five islands in the Seychelles archipelago. In the early 1900s, the species disappeared from three islands (Aride, Felicité, and Marianne) and in the 1980s disappeared from a fourth one (Praslin). Restricted to a single island (La Digue) in 1965 the population size was reduced to 28 individuals. By the year 2000, the population recovered to ∼250 individuals, relatively unassisted. In the year 2008, 23 individuals from La Digue were introduced to Denis Island and successfully established (Henriette and Laboudallon 2011). These populations continue to grow without assistance, with a current estimated species population size of 350 – 506 individuals (IUCN 2022/1).

A previous study of historical vs. modern diversity using 14 microsatellites reported a significant reduction in heterozygosity after the bottleneck (Bristol et al. 2013). Following Bristol et al. (2013), we sampled 13 historical individuals collected between 1877 to 1888 from natural history collections, and 19 modern individuals collected between 2007 and 2008 (Supplementary table 1; Fig 1A).

**Figure 1.**
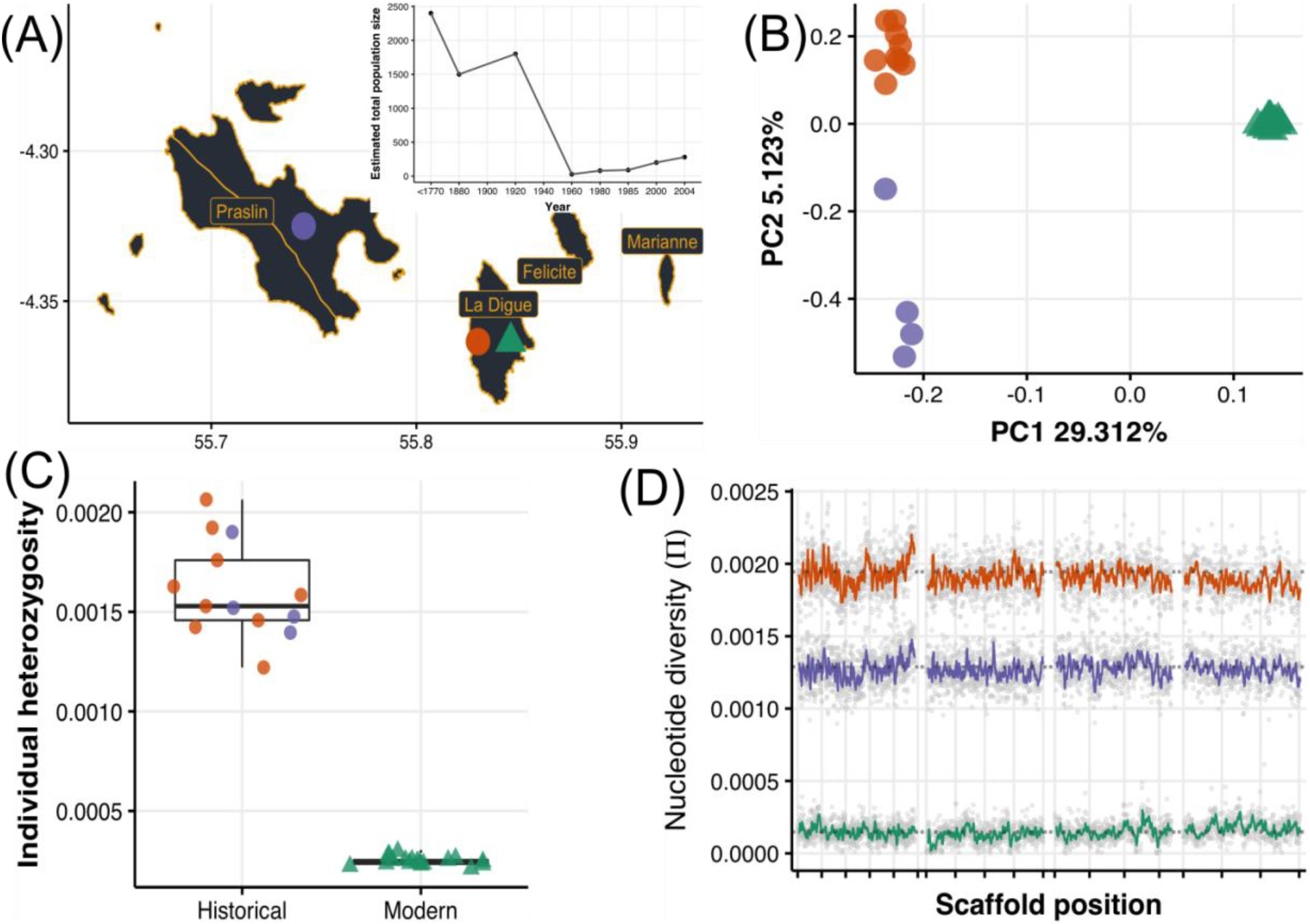
Massive loss of genome-wide diversity after population decline and despite the demographic recovery. (A) Whole-genome sequencing from historical (over 130-year-old - circles; La Digue, orange and Praslin, purple) and modern (triangles; La Digue, green) individuals. The inset shows the species’ recent demographic trajectory estimated by (Bristol et al. 2013) showing the dramatic population decline and subsequent recovery. (B) Principal component analysis of historical and modern samples. (C) On average, modern individuals have 6.4 times less observed heterozygosity than historical individuals. (D) On average, the modern population has 10.9 times less nucleotide diversity than the historical populations, and diversity was lost uniformly through the genome (here we show the longest four scaffolds; the rest of the scaffolds can be found in Fig. S6). The reason for the lower π in Praslin is that pairwise nucleotide diversity is sensitive to sample size, this effect is not seen in the individual average heterozygosity (panel C).

### Genomic libraries and sequencing

Historical DNA extractions were carried out with a modified version of Campos & Gilbert (2012) in a PCR-free clean laboratory exclusively designated for ancient DNA. Historical single-stranded sequencing libraries were prepared following the Santa Cruz Reaction protocol (Kapp et al. 2021), as modified for the DNBSEQ-G400 sequencing platform (van Grouw et al. in review) and amplified in three indexed PCR reactions. Modern DNA extractions were done with the DNAeasy commercial kit (Qiagen) following the manufacturer’s recommendations and directly submitted to BGI Copenhagen for sequencing in their DNBSEQ-G400 platform.

### Sample processing

We mapped the sequencing reads to the publicly available *de novo* sequenced reference genome of the Seychelles paradise flycatcher produced by the B10K consortium (Zhang et al. 2015), available at https://b10k.scifeon.cloud/#/b10k/Sample/S15237). We ran the automated pipeline PALEOMIX (Schubert et al. 2014) per sample for both the historical and the modern datasets. This pipeline carries out the pre-processing steps of removing adapters and collapsing mate reads, read mapping with BWA (Li and Durbin 2010), and quantifies the post-mortem damage of historical samples by running MapDamage (Ginolhac et al. 2011) We employed the *aln* algorithm for mapping historical samples and the *mem* algorithm for modern samples. Duplicates were removed with picard MarkDuplicates (Anon) and InDels were realigned with GATK (Anon).

We performed a depth-based analysis to identify and remove sex chromosome-derived scaffolds following Pečnerová et al. (2021) as their sex-biased pattern of inheritance can bias genetic diversity estimates. Considering that females are the heterogametic sex in birds (ZW), we calculated the difference in the normalized average depth between males and females, expecting that coverage will be nearly double in males relative to females for the Z chromosome, and absent in males but present in females for the W chromosome. In total, 1308 potential sex-linked scaffolds were removed from the subsequent analyses (73.4 Mb putatively Z-linked and 10.7 Mb putatively W-linked).

The small population size and population structuring can result in the sampling of closely related individuals which would inflate the estimated level of inbreeding. To avoid this, we identified and removed closely related individuals using NGSrelate2 (Hanghøj et al. 2019). We used a threshold of KING >= 0.25, R0 <= 0.1 and R1 >= 0.5 as described in Waples et al. (2019). We found only one closely related pair in the modern dataset, and we removed one of these individuals from the final dataset (Fig. S1). The pedigree metadata confirmed these two individuals had a parent-offspring relationship. The final dataset consisted of 13 historical samples (coverage: mean = 4.7 sd = 1) and 18 modern samples (coverage: mean = 9.2 sd = 0.4).

### Historical DNA biases

Historical DNA is subject to post-mortem DNA damage and contamination. This commonly leads to short sequencing reads that are error-prone and have lower quality, and samples that have low endogenous content and a low depth of coverage. We took several steps to counteract these challenges. First, we confirmed that two common features of ancient DNA datasets: reduced average sequence lengths and low coverage, did not generate reference genome mapping biases (Gopalakrishnan et al. 2022) in our historical dataset (Fig. S2). Second, we used dedicated software for low-coverage samples, ANGSD 0.921 (Korneliussen et al. 2014), to estimate genotype likelihoods and avoid directly calling genotypes. Across all ANGSD methods, we used the GATK algorithm, filtered for a base quality of 20 and a mapping quality of 30. For each population or group of samples, we computed the 1 and 99% quantiles of global depth to filter out regions with extremely low and extremely high depth. For SNP calling we inferred the major and minor alleles and used the likelihood test with a p-value threshold of 1×10<sup>−</sup>6. Third, as commonly done in ancient DNA analyses to counteract biases from post-mortem DNA damage, we removed transitions for all analyses that compared historical and modern samples.

### Population structure and genetic diversity

We performed a Principal component Analysis with PCAngsd 1.01 (Meisner and Albrechtsen 2018) with the genotype likelihoods of the joint historical-modern dataset. Next, we estimated their admixture proportions with NGSAdmix (Skotte et al. 2013), running 250 independent runs from K=2 to K=6. We evaluated the different runs using EvalAdmix (Garcia-Erill and Albrechtsen 2020) and estimated the best K using Clumpak Best K algorithm (Kopelman et al. 2015). We visualised the proportions using PONG (Behr et al. 2016).

Per-sample global heterozygosity estimates were computed directly from the site frequency spectrum (SFS) of each sample by calculating the genome-wide proportion of heterozygous genotypes. We first computed the site allele frequency (SAF) per sample in ANGSD, followed by the realSFS function to get the folded SFS assuming the reference genome as the ancestral state. We bootstrapped the SFS estimation 300 times.

To estimate the genome-wide nucleotide diversity (π) we first estimated the population-level folded SFS as done with the heterozygosity analysis but providing as input all the samples per group. We calculated per site π directly from each population’s SFS in two steps following the approach of Korneliussen et al. (2013) by dividing the pairwise Watterson theta value (Watterson 1975; Dung et al. 2019) over the effective number of sites with data (i.e., including all non-variable sites that passed the filters) per window. We computed these statistics using non-overlapping sliding windows of 50 Kb.

### Demography and runs of homozygosity

Genotypes were called with ANGSD from the genotype likelihoods as described above to identify Runs of homozygosity (ROH) in modern individuals with PLINK v1.9 (Purcell et al. 2007). SNPs not in Hardy-Weinberg equilibrium were removed and the remainder SNPs were pruned based on Linkage Disequilibrium (LD) *r*^2^ > 0.8 as implemented in Foote et al. (2021). The following parameters were used to estimate ROHs: minimum window size = 10 SNPs, minimum density per 50 kb = 1 SNP, maximum heterozygous sites per window = 5, and a maximum distance between SNPs = 1000 kb.

Analysis of recent (<100 generations) demography was performed with GONE (Santiago et al. 2020) which uses the patterns of LD to estimate recent population size changes. We used the unphased and unpruned genotypes of the modern samples as described for the ROHs and assumed a recombination rate of 3 cM/Mb with 40 replicates and default parameters.

Long-term (>5,000 generations) demography analysis was calculated with PSMC (Li and Durbin 2011) using the publicly available reference genome that was sequenced to a depth of coverage of 75x. The consensus diploid sequence was computed using SAMTOOLS and bcftools (Danecek et al. 2021). The settings for the PSMC were as follows: -N30 -t5 -r5 -p “4+30*2+4+6+10” following Nadachowska-Brzyska et al. (2015). A total of 100 independent bootstrap rounds were combined and the final plot was generated assuming a mutation rate of 4.6e-9 (as reported in the collared flycatcher; Smeds et al. 2016) and a generation time of 2 years (R Bristol unpublished data).

### Genetic load analyses

We called high-quality SNPs in each of the historical and modern individuals with bcftools (Danecek et al. 2021) retaining sites with a minimum base and mapping quality of 30, a minimum depth of 4X and a maximum of 54X, and ignoring InDels and their surrounding SNPs (5 bp). We individually annotated each filtered SNP file with SNPeff v.4.3. (Cingolani et al. 2012) using a custom database with our annotated reference genome. We classified putatively deleterious variants into (i) Low-impact variants that are likely to be not deleterious (i.e., synonymous), (ii) Moderate-impact variants that are likely to modify the protein effectiveness (i.e., missense), and (iii) High-impact variants are likely to disrupt the protein function (i.e. loss of function LoF, stop codons, splice donor variant and splice acceptor, or start codon lost) (Cingolani et al. 2012). In order to polarise our SNPs and define which were the likely ancestral and derived allelic states, we created a consensus reference by mapping the reads of two sister species; *T. cinnamomea* (estimated divergence time 4.24 MYA; (Jønsson et al. 2016; Kumar et al. 2022) and *Myiagra hebetiorwith* (11.2 MYA; (Fabre et al. 2014; Kumar et al. 2022). We randomly sampled one base with ANGSD v0.921 (Korneliussen et al. 2014) when there was no consensus across the two species.

We next compared the temporal changes in putative deleterious mutations to examine the impacts of genetic drift and purifying selection. We counted the number of homozygous derived alleles (i.e., derived counts * 2), and the number of heterozygotes. The sum of these counts represents the total number of deleterious alleles, and the number of homozygous-derived alleles represents the count of realised (expressed) load. Partially recessive deleterious mutations also contribute to the realised load. However, because the dominance coefficients (*h*) of these mutations are unknown and likely to be *h*<<0.5, we ignored this part of the realised load. We performed these calculations for moderate-, and high-impact variants. Since the historical and modern samples have different sequence quality that impacts our ability to call SNPs, we corrected these allelic counts by dividing them by the total count of synonymous sites (i.e., low-impact variants), following Kuang et al. (2020).

### Individual-based simulations

We performed individual-based forward simulations with SLiM v3.6 (Haller and Messer 2019) with a non-Wright-Fisher implementation. Absolute fitness (i.e., probability of survival) was regulated by genetic effects (see below) and the carrying capacity, which was determined with the reconstructed pre-bottleneck population size (see Results) and the known trajectory of the population decline and recovery (Bristol et al. 2013). We implemented three scenarios with different ancestral population sizes starting from the estimated ancestral population size, and population sizes that were 5 and 10 times larger (i.e. 1X, 5X or 10X). We ran a burn-in for a number of generations that was five times the population size to obtain an ancestral population in mutation-selection-drift equilibrium. We ran 100 replicates per scenario.

To confirm that our model successfully replicated the overall biology of the Seychelles paradise flycatcher, we parameterised the model with known distributions for age-based mortality probability and litter size (Currie et al. 2005). We then analysed the resulting full genealogy (with Tree sequence recording; (Haller et al. 2019)) to estimate the emerging generation time in the simulation, which matched the known generation time of ∼2 years in this species (R. Bristol unpublished data).

#### Genetics parameters

we simulated 10,000 genes of 1 Kb each distributed across 28 autosomal chromosomes, typical of a passerine genome. We used a recombination rate of 1e-4 per base position, per generation, with no recombination within genes. We use a relatively large mutation rate of 1e-7 to compensate for the small simulated genome size and ensure the accumulation of a realistic amount of genetic load in the ancestral populations.

To investigate the effect of the genetic load, we simulated deleterious mutations with selection coefficients (*s*) taken from a gamma distribution with a shape of 0.5 and scale of 0.1, plus 5% of lethal mutations, and dominance coefficient (*h*) that followed a negative relationship with *s* given by *h*=0.5*10^−13*s*^, as implemented in Kardos et al., (2021). These distributions are approximately consistent with the predicted fitness effects of deleterious variation in humans (Eyre-Walker and Keightley 2007) meta-analysis (Charlesworth and Willis 2009) and experimental approaches (Agrawal and Whitlock 2012).

To investigate the effect of adaptive potential, we simulated the additive effect of genotype values (*z*) on a polygenic trait tracking an environmental optimum (*opt*). Genotype values (*z*) were drawn from a uniform distribution ranging between -0.25 to 0.25, and with a fixed additive effect (*h*=0.5). The effect of homozygous loci was estimated as *∑z*, the effect of the heterozygous loci as *∑zh*, and the phenotype (*P*) of an individual was the sum of the homozygous and heterozygous effects. Following Falconer and Mackay (1996), we calculated the fitness effect from the deviation of the phenotype to the environmental optimum as *w =* (*P* − *opt*)^2^ and the additive genetic variation as 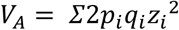. We perform an extensive parameter space exploration to test the effect of (i) the range from which genotype values (z) were drawn for the polygenic trait, and (ii) the relative proportion of mutation contributing to the adaptive trait relative to those contributing to the genetic load (Fig. S8-S10).

## Results

### Population structure and genetic diversity

Historical and modern samples (Fig 1A) showed a pattern of strong genetic differentiation, with the modern samples forming a homogenous group (PC1; 29% explained variance), and the historical samples differentiated between islands (PC2 5% explained variance) (Fig 1B). The rest of the PCs mostly account for the variation within the historical populations (Fig S3). The admixture analysis assuming three genetic groups (K=3) reflects the strong differentiation between historical and modern individuals and the geographical structure within the historical individuals (Fig S4). Higher Ks yielded no clear signal of co-ancestry between the historical and modern La Digue individuals. This failure to retrieve a historical component in the modern samples is likely due to strong genetic drift changing the allele frequencies in the modern population (Ebenesersdóttir et al. 2018).

On average, the global individual heterozygosity of modern individuals (La Digue: mean=0.00024, sd=0.00002) was 6.4 times lower than that of the historical individuals (La Digue: mean=0.00162, sd=0.0003 and Praslin: mean=0.00157, sd=0.0002) (Fig. 1C). The genomic sliding-window analysis of population pairwise nucleotide diversity shows that the loss with this metric was 10.9-fold, and that genetic diversity was lost similarly throughout the entire genome (Fig. 1D). Similar comparisons in different bird species have reported smaller losses in nucleotide diversity: the crested ibis and the Chatham Island black robin with a 1.8 fold loss (Feng et al. 2019; von Seth et al. 2022), or in heterozygosity levels: the New Zealand Saddleback, 4.16 fold (Taylor et al. 2007); the Mangrove Finch: 1.32 fold (Lawson et al. 2017); Greater Prairie Chicken: 1.26 fold (Bellinger et al. 2003) (Supplementary Table 2). The resulting extremely low genetic diversity in the modern population of the Seychelles paradise flycatcher is considerably lower compared to other threatened bird species (Fig. S5). Our results highlight how even when a population has recovered demographically, it can still be a long way away from recovering in terms of genetic diversity.

### Demography and runs of homozygosity

The modern La Digue population has an overall absence of long (>5Mb) Runs of Homozygosity (ROHs) (Fig. 2A). Long ROHs would be expected if closely related individuals mated with each other within the last 10 generations (Fig 2B; Fig. S7). Hence, the lack of long ROH suggests an absence of recent inbreeding in our data (F_ROH_ < 0.01; Fig. 2B). Instead, the vast majority of ROHs are between 1-2 Mb long (Fig 2A). This class of ROH length is expected to have been formed 10-20 generations ago (Fig. S7), which is consistent with historical inbreeding (F_ROH_ = 0.2-0.4; Fig. 2B) around the year 1974, and it is likely to be a product of the bottleneck that started in the mid-1960s (Fig 2B).

**Figure 2.**
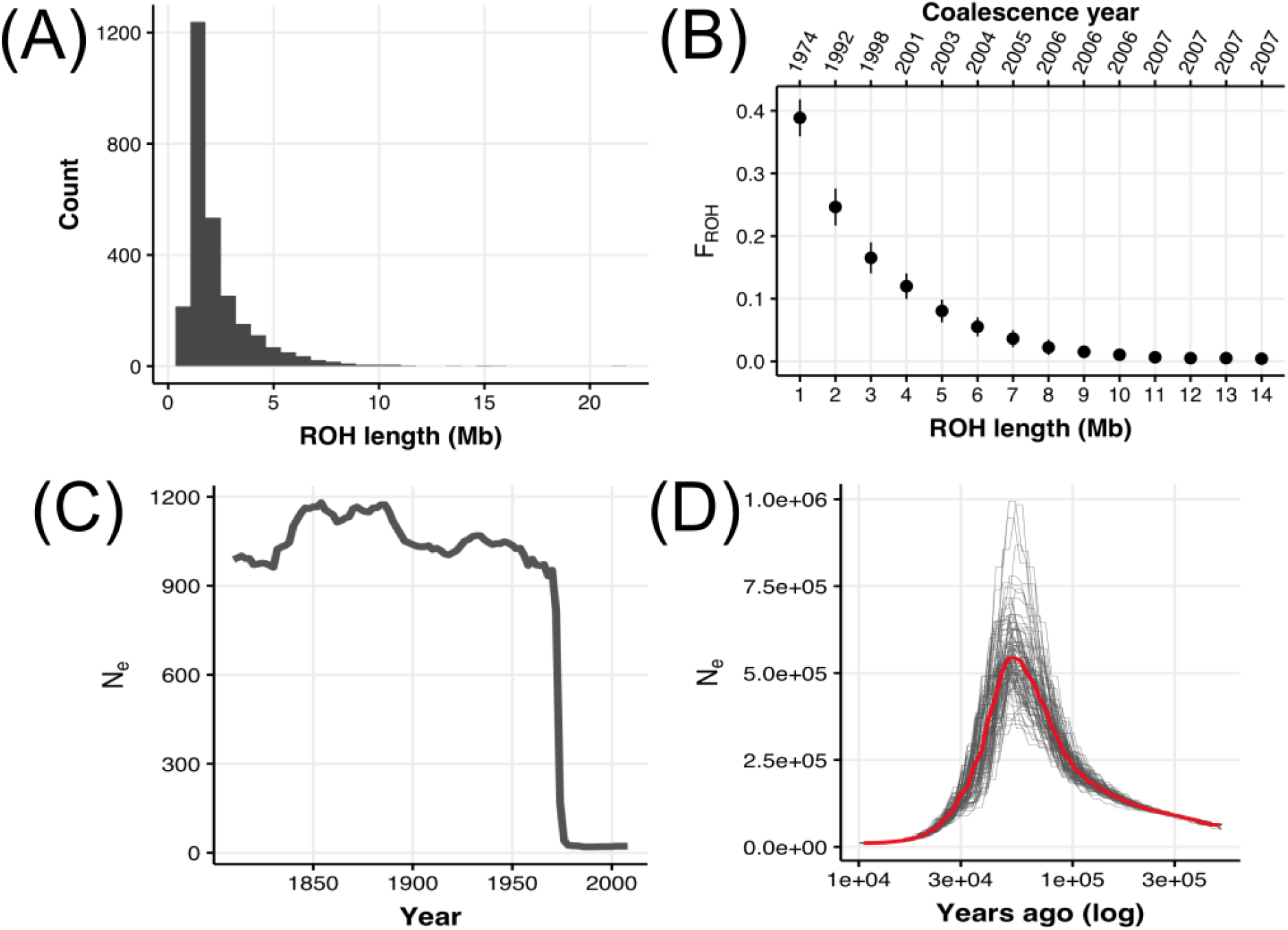
Demographic reconstruction of population decline. (A) Runs of homozygosity (ROH) length distribution across all modern individuals. (B) Inbreeding coefficient estimated for different classes of ROH lengths (F_ROH_). The year represents the estimated time at which a category of ROH length was formed assuming a recombination rate of 3 cM/Mb and using the formula L = 100/2t cM from (Thompson 2013), where *L* is the ROH length and *cM* is the recombination rate, to obtain *t* the time of ROH coalescence in generations. Generations ago were converted assuming a generation time of 2 years from the time of sampling. (C) Reconstruction of the recent demography (last 100 generations) from Linkage Disequilibrium using GONE (Santiago et al. 2020)assuming a recombination rate of 3 cM/Mb. Generations ago were converted assuming a generation time of 2 years from the time of sampling (2010). (D) Reconstruction of the ancient demography (<10,000 years ago or 5,000 generations ago) from genetic coalescence using PSMC (Li and Durbin 2011) assuming a mutation rate of 2.3e-9 and a generation time of 2 years.

The reconstructed recent demographic history (within the last 100 generations) with GONE (Santiago et al. 2020) also recovered a clear signature of the bottleneck by registering a dramatic drop in the N_e_ around the year 1975 (∼17 generations ago; Fig. 2C), in full agreement with the F_ROH_ evidence. Before the bottleneck, the ancestral population had a steady and relatively small N_e_, as expected from a small insular species. Our reconstruction of the ancient and recent demographic history of the Seychelles paradise flycatcher is consistent with a scenario that prior to the recent bottleneck of 1964, the species had a long-term small N_e_ for the last ∼5,000 generations or ∼10,000 years. It is important to note that the deep demographic history reconstruction with PSMC carries some uncertainty. The maximum N_e_=530,635 estimated at ∼55,000 years substantially exceeds the current carrying capacity of the Seychelles archipelago. However, past the sea levels were highly dynamic, connecting and disconnecting islands in the archipelago on at least 14 separate occasions (Ryan et al. 2009; Warren et al. 2010; Ali 2018). Thus, it is possible that the ancient population could have been much larger at times of increased island connectivity. Seychelles’ landmass is estimated to have been up to 180 times its present size, and gene flow may have been facilitated by islands in the western Indian Ocean that could have acted as stepping-stones between landmasses during the Pliocene and Pleistocene (Warren et al. 2010; Cheke and Hume) This geological signature has been seen in other Seychelles taxa (Groombridge et al. 2002; Rocha et al. 2013; Labisko et al. 2022). However, the large ancestral N_e_ can also be an artefact of population structure, selection and admixture, all of which are known to introduce biases to coalescent demographic reconstruction (Mazet et al. 2016; Johri et al. 2021; Boitard et al. 2022). For example, if island populations were reproductively separated at some point, PSMC estimates would be inflated as alleles would not coalesce. Irrespective of the uncertainty of ancient N_e_ estimates, we can be confident that the relatively-recent genetic lineage remained small for at least 5,000 generations (10,000 years). This is consistent with a history of long-term small N_e_ which would have led to the purging of the genetic load, making the population more resilient against contemporary inbreeding.

### Genetic load analyses

Consistent with theoretical expectations (Hedrick and Garcia-Dorado 2016; Bertorelle et al. 2022; van Oosterhout et al. 2022), and empirical observations in other taxa (Dussex et al. 2021; Grossen et al. 2020; Khan et al. 2021; Kleinman-Ruiz et al. 2022; Kleinman-Ruiz et al. 2022), we observed a lower proportion of deleterious mutations in the modern population relative to the historical population (Fig. 3A). This pattern is expected given the massive loss of genetic diversity as many deleterious alleles were likely lost during the bottleneck due to drift. We also found a reduction in the count of heterozygous deleterious alleles in the modern population (Fig. 3B), consistent with the observed genome-wide loss in heterozygosity, and with heterozygous deleterious mutations being unmasked into their homozygous state (i.e., genetic load conversion; Bertorelle et al. 2022; Mathur and DeWoody 2021). In line with the expectation of genetic load purging, we found that the proportion of derived alleles (i.e., realised load) classified as high-impact (i.e., probably highly deleterious) is lower in the modern population (Fig. 3C). In contrast, mutations classified as mildly deleterious remained the same (Fig. 3D). Due to their lower selection coefficients, these mildly deleterious mutations would have behaved like near-neutral variants during the bottleneck, which explains why their ratio (Mild/Synonymous) did not change.

**Figure 3.**
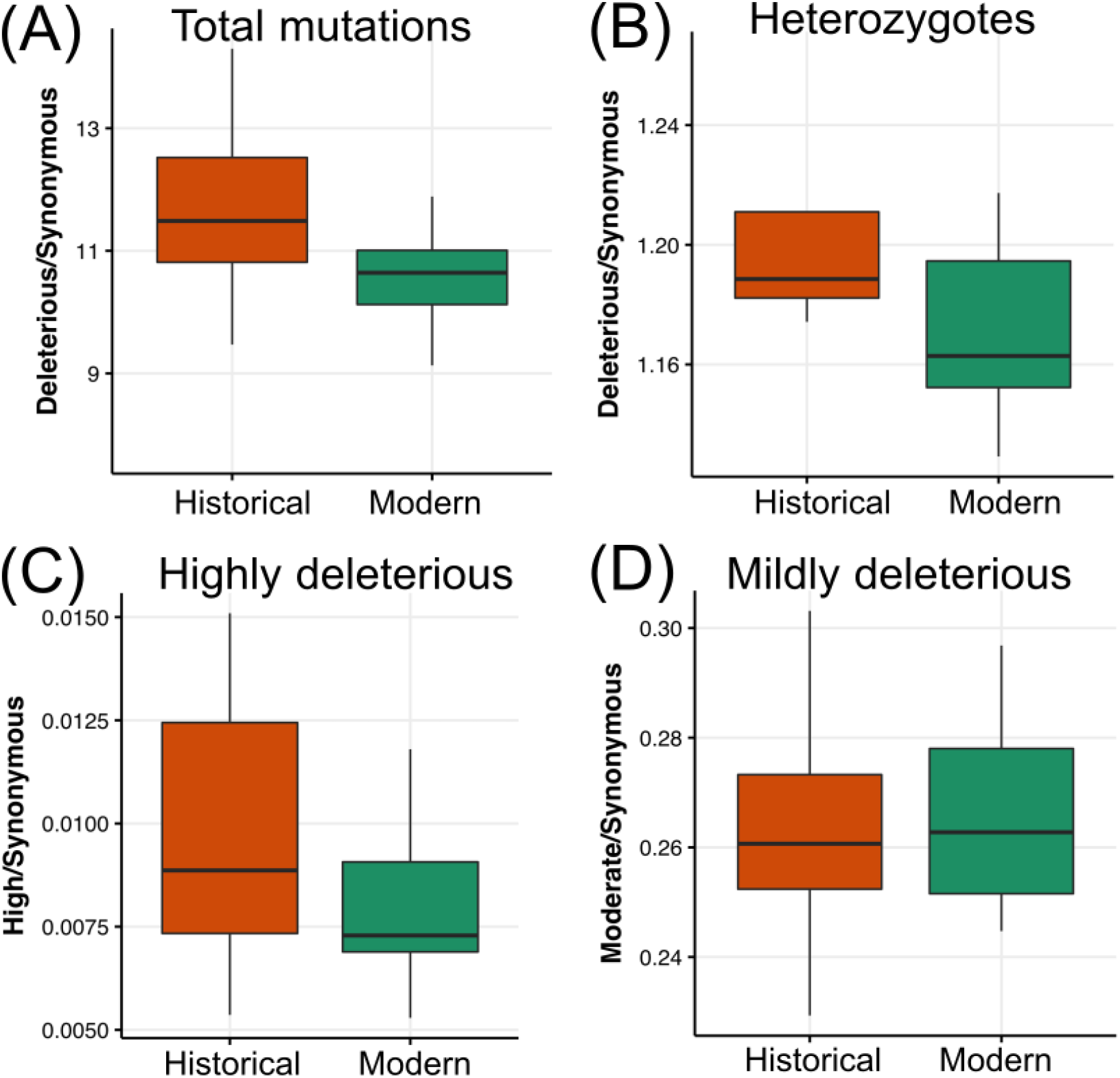
Purging and accumulation of genetic load. We compared allelic counts between the two time points in La Digue historical (orange) and La Digue modern (green) populations. To avoid biases from the differential ability to call alleles in the high-quality modern samples relative to the degraded historical samples, counts were divided by the count of synonymous alleles (classified as low). (A) Total count of all deleterious alleles (high + moderate). (B) Count of heterozygous deleterious alleles (high + moderate). (C) Count of high-impact derived homozygous deleterious alleles. (D) Count of moderate-impact derived homozygous deleterious alleles.

### Individual-based simulations

We assessed how different types of genomic variation (deleterious variation and adaptive variation) respond to the population decline and recovery in the species by simulating historic populations with small (1X), medium (5X), and large (10X) ancestral population size (Fig. 4A). The total ancestral deleterious variation (i.e., genetic load = sum of masked load plus realised load) scales positively with population size (Fig. 4B). Historically, most deleterious variation is in the form of masked load (Fig. 4C) (i.e., these mutations do not reduce fitness), and only a small proportion is part of the realised load (Fig. 4D) (i.e., mutations that reduce fitness). During the population size collapse, there is a marginal reduction of the genetic load (Fig. 4B) as many rare, low-frequency variants are lost due to drift.

**Figure 4.**
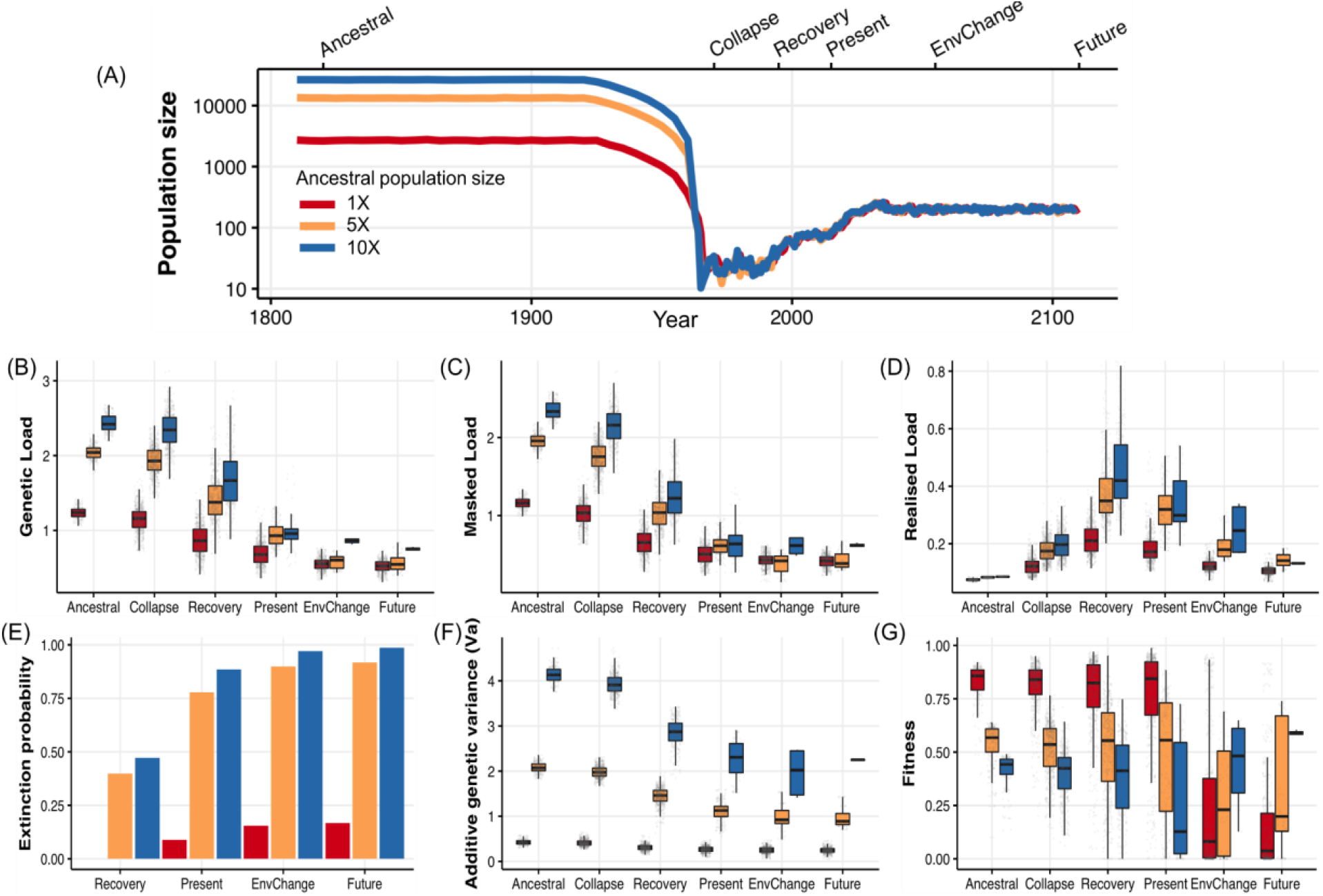
Forward simulations of deleterious and adaptive variation. (A) Alternative simulated demographic trajectories. The 1X trend (red) represents the known ancestral size of the Seychelles paradise flycatcher based on the reconstruction of the recent demography with GONE (Fig. 2C). The alternative scenarios represent medium (5X, yellow) and large (10X, blue) ancestral population sizes. The trajectory was divided into six stages: Ancestral (years 1810-1815), Collapse (1965-1970), Recovery (1990-1995), Present (2010-2015), Environmental change (2050-2055), Future (2095-2100). During the environmental shift, the quantitative trait optimum value changed from 0.2 to 1.2. (B) Genetic load. (C) Masked load. (D) Realised load. Genetic load is expressed in lethal equivalents (see (Bertorelle et al. 2022). The reduction in fitness (w) due to the expression of unconditionally deleterious mutations (i.e., inbreeding depression) is a function of the realised load: w = e^-Realised Load^. (E) Extinction probability per scenario (the number of surviving replicates divided by the total number of replicates). (F) Fitness effect conferred by the quantitative trait. (G) Additive genetic variance in the quantitative trait.

On the flip side, during the bottleneck, some of the masked load is converted into realised load by inbreeding (Fig. 4C and D), and this results in a loss of fitness (i.e., inbreeding depression). Two processes are at play here. First, whilst most deleterious variants are lost, genetic drift increases the frequency of a small number of deleterious mutations. Given their now elevated frequency, these deleterious mutations are more likely to be found in homozygous genotypes. Second, the bottleneck increases the probability of mating between closely related individuals. By increasing homozygosity, both genetic drift and inbreeding convert the masked load into a realised load. Figure 4D illustrates this in computer simulations. During the population size collapse, the realised load of the large ancestral population is increased to around 0.2 lethal equivalents, which equates to a fitness w = e^-0.2^ = 0.82. For the smallest ancestral population size, population size collapse increases the realised load to circa 0.1 lethal equivalents, which equates to w = e^-0.1^ = 0.90. In other words, individuals in large ancestral populations suffer from more severe inbreeding depression during population size collapse than individuals derived from historically small populations.

Remarkably, we observed the most significant changes in the magnitude and composition of the genetic load during population recovery (Fig. 4 B-D). This highlights an important concept; the full effects of genetic drift and inbreeding are only felt after the bottleneck, and they continue to impact fitness and population viability despite the population-size recovery. This is known as the “drift debt” (Gilroy et al. 2017). In particular, the realised load peaked at over 0.4 lethal equivalents with the largest ancestral population size during recovery. Severe inbreeding depression during this stage would have reduced the fitness of individuals markedly, w = e^-0.4^ = 0.67 (i.e., 33% individual fitness loss in average). Figure 4D shows that the worst affected individuals express a realised load of 0.8 lethal equivalents, which means that their fitness would be less than half that of their pre-bottleneck ancestors. In contrast, the smallest ancestral population (i.e., the simulations most similar to the Seychelles paradise flycatcher) suffer much less inbreeding depression during their recovery. An average individual is expected to express 0.2 lethal equivalents, a ∼18% reduction in fitness. This explains why the small population has a much lower extinction risk (Fig. 4E), and why the Seychelles paradise flycatcher may have avoided extinction.

Unlike unconditionally deleterious mutations that always reduce fitness when expressed, additive genetic variants can either increase or decrease fitness depending on the genetic background and the environment (Charlesworth 2013a; Charlesworth 2013b). Take for example an additive genetic variant that increases the trait value. If this variant is present in an individual with a suboptimally large trait, it would reduce fitness (i.e., it would be negatively selected). Conversely, that same variant would be beneficial in an individual with a suboptimally small trait. Figure 4F shows that the amount of additive genetic variation (V_a_) increases with population size. In a stable environment, ancestrally larger populations have on average lower fitness (Fig 4G). This is because they contain more segregating variants which can produce more extreme (i.e., suboptimal) phenotypes (Charlesworth 2013b). However, their larger V_a_ gives them a wider phenotypic breath and a greater adaptive potential when environmental conditions change. As expected, population size collapse reduces V_a_, but as with the genetic load, the effect on quantitative genetic variation is most pronounced during recovery (Fig. 4F). Remarkably, the loss in V_a_ results in the most pronounced fitness loss in large ancestral populations. However, after environmental change, recovered populations derived from large ancestral populations can better match the new environmental optimum. Their superior adaptive potential ensures that such populations have a higher fitness during environmental change in the future (Fig. 4H).

## Discussion

We analysed the whole genome sequence data of a threatened species that suffered a population decline to 28 individuals, the Seychelles paradise flycatcher (*Terpsiphone corvina*), comparing the level of genomic erosion between 13 historic (> 130-years-old) and 18 modern birds. In addition, we conducted computer simulations, to assess the effect of population decline and recovery on the genetic load and adaptive evolutionary potential to assess its long-term population viability. We uncovered a 10-fold loss of genetic diversity, reflecting severe genetic drift during population size decline that continues to act despite their recovery. We also found evidence of historical inbreeding at the time of their decline, but no evidence of recent inbreeding, a testament to their successful population recovery. Demographic reconstructions suggest that prior to its recent population decline, the Seychelles paradise flycatcher sustained a small effective population size (N_e_) for thousands of generations. Our genomic simulations suggest that this reduced the amount of genetic load in the ancestral population, resulting in only mild inbreeding depression during its collapse. In other words, the long-term small N_e_ of this species may have allowed for its (unassisted) demographic recovery and helped avoid extinction. However, the species has not recovered its genetic diversity, and the mean fitness of individuals is predicted to be lower than that of their ancestors. Our simulation also indicates that the loss of genetic diversity has likely reduced their adaptive potential, and this could jeopardise the species’ long-term viability when faced with environmental change. Our analysis illustrate the power historical vs. modern comparisons, in combination with analyses of genomic erosion and simulations to assess the effect of population decline and recovery on population viability (Díez-del-Molino et al. 2018; Feng et al. 2019; Dussex et al. 2021; Sánchez-Barreiro et al. 2021). Importantly, we showcase how to use this integrative approach to inform conservation assessments (Jensen et al. 2022; van Oosterhout et al. 2022).

### Historical inbreeding, but not recent inbreeding

Remarkably, we did not find evidence of long runs of the homozygosity (ROH), which are typically observed in recently bottlenecked species such as the crested Ibis (Feng et al. 2019), the alpine ibex (Grossen et al. 2020), the white rhinoceros (Sánchez-Barreiro et al. 2021), and different horse breeds (Grilz-Seger et al. 2018). Instead, we found that the most common category of ROH was between 1-2 Mb long. ROHs are formed when very closely related individuals mate (i.e., consanguineous mating or inbreeding). The inbred offspring inherit identical segments of DNA, which show up as ROH across the genome. Later, if this offspring mates with unrelated individuals the ROHs can be “broken down” by recombination and they become shorter. Thus, the distribution of ROH size reflects an inbreeding timeline. Our results suggest that the population decline imposed severe inbreeding, and that at the time of the bottleneck individuals had ∼40% of their genomes contained in ROHs (F_ROH_=0.4; Fig. 2B). Upon demographic recovery, ROHs were broken down by recombination, leaving this signature of relatively shorter ROHs (1-2 Mb), consistent with historical inbreeding (∼45 years ago). On the other hand, the lack of long ROHs indicated the absence of recent inbreeding (F_ROH_<0.01 in the last decade; Fig. 2B), meaning that the demographic recovery allowed the Seychelles paradise flycatcher to avoid consanguineous mating.

### Genetic load, adaptive potential, and extinction risk

Although population bottlenecks reduce the number of segregating sites with deleterious mutations, they also increase the frequency of deleterious variants at some loci. This elevated frequency increases the level of homozygosity, which increases the realised load and leads to inbreeding depression (van Oosterhout et al. 2022). In turn, this allows for purifying selection to purge some of the realised load, effectively reducing the genetic load. Consistent with these theoretical expectations, we observed a pattern of purging after the bottleneck in the Seychelles paradise flycatcher. Purging was only observed, however, for variants classified as highly-deleterious. These variants have the strongest fitness effects and are thus most effectively removed by selection. Derived alleles of mildly deleterious variants were not reduced in the modern population. This is notable, given that genetic drift and inbreeding during and after the population decline must have exposed many of these deleterious variants to selection. For this class of variants, drift overwhelmed selection, resulting in the accumulation of some mildly deleterious variation and a reduction in fitness.

It is important to note, however, that scoring deleterious alleles in the historical samples is challenging and error-prone given the low depth of coverage in our data. While we used specialised software for low-coverage data, removed transitions, and counted derived alleles relative to the count of synonymous sites to reduce this potential bias (Kuang et al. 2020), these results should be taken with caution.

Small-island species with long-term small N_e_ accumulate less masked load compared to mainland species with large ancestral N_e_. This could make small-island species more resilient to the effects of population decline and inbreeding depression. Our reconstruction of the recent demographic history of the Seychelles paradise flycatcher is consistent with a scenario that prior to the recent bottleneck of 1964, the species had a long-term small N_e_ for the last ∼5,000 generations. During population decline, inbreeding and genetic drift convert the masked load into the realised load (e.g., Mathur and DeWoody 2021; Smeds and Ellegren 2022). Natural selection removes some of this expressed deleterious variation (i.e., genetic load purging). However, a portion of the converted load escapes selection and persists as realised load, reducing population viability (Grossen et al. 2020; van Oosterhout et al. 2022). Given that historically large populations possess a high masked load, they are particularly prone to the detrimental effects of load conversion during population size collapse (Mathur and DeWoody 2021; van Oosterhout et al. 2022). Conversely, the Seychelles paradise flycatcher was particularly resilient to inbreeding depression and this likely played a role in their successful (unassisted) demographic recovery.

The Seychelles paradise flycatcher population size has been steadily increasing in the past 20 years. Nonetheless, even after the apparent demographic recovery, the modern population possesses a very low genetic diversity. This is of concern because genome-wide diversity is an important predictor of population fitness and adaptive potential (Hansson and Westerberg 2002; Fagan and Holmes 2006; Willi et al. 2006; Harrisson et al. 2014; Willoughby et al. 2015; Kardos et al. 2021; Willi et al. 2022). Our computer simulations show that this may lead to reduced adaptive response during environmental change. In turn, and opposite to the prediction for the genetic load, this could elevate its extinction risk compared to populations with a larger ancestral N_e_ (Lande and Shannon 1996; Willi et al. 2006).

### The role of genomics in species conservation assessments

The incorporation of genetic information into assessments of conservation status and policy remains inadequate (Hoban et al. 2020; Laikre et al. 2020). Here, we show the impact of genomic erosion in the Seychelles paradise flycatcher, a species that has made a successful demographic recovery that resulted in its downlisting in the Red List. Our findings suggest that its ancestrally small N_e_ might have conferred resilience to inbreeding that initially eases demographic recovery. However, it may also compromise its adaptive potential, particularly during environmental change. Moreover, it is important to note that the purging of their ancestral genetic load happened (naturally) over thousands of generations. In addition, the chronic reduction in fitness caused by an elevated realised load is likely to put the species at increased risk of extinction.This might be of particular relevance to other island endemic species, which are, for example, characterised by reduced immune function, partially due to their low N_e_ (Barthe et al. 2022). Accordingly, low genetic diversity can make species more prone to emerging infectious diseases, which is a substantial risk given the high rates of new colonisations and invasive species in islands (Sax and Gaines 2008; Lockwood et al. 2009). Our work demonstrates the power of direct comparisons between historical and modern whole genomes to reconstruct the temporal dynamics of diversity, demography and inbreeding, and the importance of combining these insights with simulations to inform conservation. We argue that the downlisting of the IUCN Red List status may sometimes be premature and requires assessing the risks posed by genomic erosion. A promising way forward to achieve this is incorporating the analysis of genomic erosion in population viability analysis (PVA) with computer simulation to leverage the full power of genomic and ecological/demographic data.

## Acknowledgments

We are grateful to curators Hein van Grouw from the Natural History Museum, Tring, UK and Michael Brooke from the Museum of Zoology, University of Cambridge, UK. Darwin Initiative grant 15/009 to JG facilitated blood-sampling of the modern population. We are grateful to members of the B10K consortium for their work on the reference genome and Ester Milesi for help processing samples. We thank Shyam Gopalakrishnan for advice on the reference bias analyses. HEM was funded by the European Union’s Horizon 2020 Research and Innovation Programme under a Marie Sklodowska-Curie grant (840519). MTPG acknowledges DNRF143 award for funding. CvO was funded by the Earth and Life Systems Alliance (ELSA). GF was supported by UNAM-DGECI Iniciación a la Investigación.

## Data availability

The reference genome can be found at https://b10k.scifeon.cloud/#/b10k/Sample/S15237. The raw sequencing reads have been deposited in the Sequence Read Archive under the accession number XXX-XXX. Scripts for the SLiM simulations can be found at https://github.com/hmoral/SPF

## Author contributions

HEM, MTPG and JG conceived the study. GF generated the data. GF and HEM performed the analysis. CvO and JG provided insights to interpret the result. JG and RMB provided samples and insights about the study species. HEM and MTPG provided resources. GZ and SF developed the reference genome. HEM wrote the manuscript with the assistance of GF, CvO and MTPG. All authors provided feedback and approved the final version.

## References

Agrawal AF, Whitlock MC. 2012. Mutation Load: The Fitness of Individuals in Populations Where Deleterious Alleles Are Abundant. Annu. Rev. Ecol. Evol. Syst. 43:115–135.

Ali JR. 2018. Islands as biological substrates: Continental. J. Biogeogr. 45:1003–1018.

Almond REA, Grooten M, Peterson T. 2022. Living Planet Report 2020-Bending the curve of biodiversity loss. Available from: http://pure.iiasa.ac.at/id/eprint/16870/1/ENGLISH-FULL.pdf

Anon. Picard. Available from: http://broadinstitute.github.io/picard/

Anon. GATK. Available from: https://gatk.broadinstitute.org/hc/en-us

Barthe M, Doutrelant C, Covas R, Melo M, Illera JC, Tilak M-K, Colombier C, Leroy T, Loiseau C, Nabholz B. 2022. Evolution of immune genes in island birds: reduction in population sizes can explain island syndrome. Peer Community Journal [Internet] 2. Available from: https://peercommunityjournal.org/articles/10.24072/pcjournal.186/

Behr AA, Liu KZ, Liu-Fang G, Nakka P, Ramachandran S. 2016. pong: fast analysis and visualization of latent clusters in population genetic data. Bioinformatics 32:2817–2823.

Bellinger MR, Johnson JA, Toepfer J, Dunn P. 2003. Loss of genetic variation in greater prairie chickens following a population bottleneck in Wisconsin, U.s.a. Conserv. Biol. 17:717–724.

Bertorelle G, Raffini F, Bosse M, Bortoluzzi C, Iannucci A, Trucchi E, Morales HE, van Oosterhout C. 2022. Genetic load: genomic estimates and applications in non-model animals. Nat. Rev. Genet. 23:492–503.

Boitard S, Arredondo A, Chikhi L, Mazet O. 2022. Heterogeneity in effective size across the genome: effects on the inverse instantaneous coalescence rate (IICR) and implications for demographic inference under linked selection. Genetics [Internet] 220. Available from: http://dx.doi.org/10.1093/genetics/iyac008

Bozzuto C, Biebach I, Muff S, Ives AR, Keller LF. 2019. Inbreeding reduces long-term growth of Alpine ibex populations. Nature Ecology & Evolution [Internet] 3:1359–1364. Available from: http://dx.doi.org/10.1038/s41559-019-0968-1

Bristol RM, Tucker R, Dawson DA, Horsburgh G, Prys-Jones RP, Frantz AC, Krupa A, Shah NJ, Burke T, Groombridge JJ. 2013. Comparison of historical bottleneck effects and genetic consequences of re-introduction in a critically endangered island passerine. Mol. Ecol. 22:4644–4662.

Charlesworth B. 2013a. Why we are not dead one hundred times over. Evolution 67:3354–3361.

Charlesworth B. 2013b. Stabilizing selection, purifying selection, and mutational bias in finite populations. Genetics 194:955–971.

Charlesworth D, Willis JH. 2009. The genetics of inbreeding depression. Nat. Rev. Genet. 10:783–796.

Cheke AS, Hume JP. Lost land of the Dodo: the ecological history of the Mascarene Islands. T and AD Poyser, London.

Cingolani P, Platts A, Wang LL, Coon M, Nguyen T, Wang L, Land SJ, Lu X, Ruden DM. 2012. A program for annotating and predicting the effects of single nucleotide polymorphisms, SnpEff: SNPs in the genome of Drosophila melanogaster strain w1118; iso-2; iso-3. Fly 6:80–92.

Currie D, Bristol R, Millett J, Shah NJ. 2005. Demography of the Seychelles Black Paradise-flycatcher: considerations for conservation and reintroduction. Ostrich 76:104–110.

Danecek P, Bonfield JK, Liddle J, Marshall J, Ohan V, Pollard MO, Whitwham A, Keane T, McCarthy SA, Davies RM, et al. 2021. Twelve years of SAMtools and BCFtools. Gigascience 10: giab008.

Díez-del-Molino D, Sánchez-Barreiro F, Barnes I, Gilbert MTP, Dalén L. 2018. Quantifying Temporal Genomic Erosion in Endangered Species. Trends Ecol. Evol. 33:176–185.

Dung SK, López A, Barragan EL, Reyes R-J, Thu R, Castellanos E, Catalan F, Huerta-Sánchez E, Rohlfs RV. 2019. Illuminating Women’s Hidden Contribution to Historical Theoretical Population Genetics. Genetics 211:363–366.

Dussex N, van der Valk T, Morales HE, Wheat CW, Díez-del-Molino D, von Seth J, Foster Y, Kutschera VE, Guschanski K, Rhie A, et al. 2021. Population genomics of the critically endangered kakapo. Cell Genomics 1:100002.

Ebenesersdóttir SS, Sandoval-Velasco M, Gunnarsdóttir ED, Jagadeesan A, Guðmundsdóttir VB, Thordardóttir EL, Einarsdóttir MS, Moore KHS, Sigurðsson Á, Magnúsdóttir DN, et al. 2018. Ancient genomes from Iceland reveal the making of a human population. Science 360:1028–1032.

Eyre-Walker A, Keightley PD. 2007. The distribution of fitness effects of new mutations. Nat. Rev. Genet. 8:610–618.

Fabre P-H, Moltensen M, Fjeldså J, Irestedt M, Lessard J-P, Jønsson KA. 2014. Multiple waves of colonization by monarch flycatchers (Myiagra, Monarchidae) across the Indo-Pacific and their implications for coexistence and speciation. J. Biogeogr. 41:274–286.

Fagan WF, Holmes EE. 2006. Quantifying the extinction vortex. Ecol. Lett. 9:51–60.

Falconer DS, Mackay TFC. Introduction to quantitative genetics 4th edition. Harlow, UK: Longmans.

Feng S, Fang Q, Barnett R, Li C, Han S, Kuhlwilm M, Zhou L, Pan H, Deng Y, Chen G, et al. 2019. The Genomic Footprints of the Fall and Recovery of the Crested Ibis. Curr. Biol. 29:340–349.e7.

Foote AD, Hooper R, Alexander A, Baird RW, Baker CS, Ballance L, Barlow J, Brownlow A, Collins T, Constantine R, et al. 2021. Runs of homozygosity in killer whale genomes provide a global record of demographic histories. Mol. Ecol. 30:6162–6177.

Forester BR, Beever EA, Darst C, Szymanski J, Funk WC. 2022. Linking evolutionary potential to extinction risk: applications and future directions. Front. Ecol. Environ. 20:507–515.

Garcia-Erill G, Albrechtsen A. 2020. Evaluation of model fit of inferred admixture proportions. Mol. Ecol. Resour. 20:936–949.

Gilroy DL, Phillips KP, Richardson DS, van Oosterhout C. 2017. Toll-like receptor variation in the bottlenecked population of the Seychelles warbler: computer simulations see the “ghost of selection past” and quantify the “drift debt.” J. Evol. Biol. 30:1276–1287.

Ginolhac A, Rasmussen M, Gilbert MTP, Willerslev E, Orlando L. 2011. mapDamage: testing for damage patterns in ancient DNA sequences. Bioinformatics 27:2153–2155.

Gopalakrishnan S, Ebenesersdóttir SS, Lundstrøm IKC, Turner-Walker G, Moore KHS, Luisi P, Margaryan A, Martin MD, Ellegaard MR, Magnússon ÓÞ, et al. 2022. The population genomic legacy of the second plague pandemic. Curr. Biol. [Internet]. Available from: http://dx.doi.org/10.1016/j.cub.2022.09.023

Grilz-Seger G, Mesarič M, Cotman M, Neuditschko M, Druml T, Brem G. 2018. Runs of Homozygosity and Population History of Three Horse Breeds With Small Population Size. J. Equine Vet. Sci. 71:27–34.

Groombridge JJ, Jones CG, Bayes MK, van Zyl AJ, Carrillo J, Nichols RA, Bruford MW. 2002. A molecular phylogeny of African kestrels with reference to divergence across the Indian Ocean. Mol. Phylogenet. Evol. 25:267–277.

Grossen C, Guillaume F, Keller LF, Croll D. 2020. Purging of highly deleterious mutations through severe bottlenecks in Alpine ibex. Nat. Commun. 11:1001.

Haller BC, Galloway J, Kelleher J, Messer PW, Ralph PL. 2019. Tree-sequence recording in SLiM opens new horizons for forward-time simulation of whole genomes. Mol. Ecol. Resour. 19:552–566.

Haller BC, Messer PW. 2019. SLiM 3: Forward Genetic Simulations Beyond the Wright–Fisher Model. Mol. Biol. Evol. 36:632–637.

Hanghøj K, Moltke I, Andersen PA, Manica A, Korneliussen TS. 2019. Fast and accurate relatedness estimation from high-throughput sequencing data in the presence of inbreeding. Gigascience [Internet] 8. Available from: http://dx.doi.org/10.1093/gigascience/giz034

Hansson B, Westerberg L. 2002. On the correlation between heterozygosity and fitness in natural populations. Mol. Ecol. 11:2467–2474.

Harrisson KA, Pavlova A, Telonis-Scott M, Sunnucks P. 2014. Using genomics to characterize evolutionary potential for conservation of wild populations. Evol. Appl. 7:1008–1025.

Hedrick PW, Garcia-Dorado A. 2016. Understanding Inbreeding Depression, Purging, and Genetic Rescue. Trends Ecol. Evol. 31:940–952.

Henriette E, Laboudallon V. 2011. Seychelles paradise flycatcher conservation introduction: population assessment on Denis Island, Seychelles. Unpublished report.

Hoban S, Bruford M, D’Urban Jackson J, Lopes-Fernandes M, Heuertz M, Hohenlohe PA, Paz-Vinas I, Sjögren-Gulve P, Segelbacher G, Vernesi C, et al. 2020. Genetic diversity targets and indicators in the CBD post-2020 Global Biodiversity Framework must be improved. Biol. Conserv. 248:108654.

IUCN. 2022. IUCN Red List of Threatened Species. IUCN Red List of Threatened Species [Internet]. Available from: https://www.iucnredlist.org

Jensen EL, Díez-Del-Molino D, Gilbert MTP, Bertola LD, Borges F, Cubric-Curik V, de Navascués M, Frandsen P, Heuertz M, Hvilsom C, et al. 2022. Ancient and historical DNA in conservation policy. Trends Ecol. Evol. 37:420–429.

Johri P, Riall K, Becher H, Excoffier L, Charlesworth B, Jensen JD. 2021. The Impact of Purifying and Background Selection on the Inference of Population History: Problems and Prospects. Mol. Biol. Evol. 38:2986–3003.

Jønsson KA, Fabre P-H, Kennedy JD, Holt BG, Borregaard MK, Rahbek C, Fjeldså J. 2016. A supermatrix phylogeny of corvoid passerine birds (Aves: Corvides). Molecular Phylogenetics and Evolution [Internet] 94:87–94. Available from: http://dx.doi.org/10.1016/j.ympev.2015.08.020

Kapp JD, Green RE, Shapiro B. 2021. A Fast and Efficient Single-stranded Genomic Library Preparation Method Optimized for Ancient DNA. J. Hered. [Internet] 112. Available from: https://pubmed.ncbi.nlm.nih.gov/33768239/

Kardos M, Armstrong EE, Fitzpatrick SW, Hauser S, Hedrick PW, Miller JM, Tallmon DA, Funk WC. 2021. The crucial role of genome-wide genetic variation in conservation. Proc. Natl. Acad. Sci. U. S. A. [Internet] 118. Available from: http://dx.doi.org/10.1073/pnas.2104642118

Khan, A., Patel, K., Shukla, H., Viswanathan, A., van der Valk, T., Borthakur, U., … & Ramakrishnan, U. (2021). Genomic evidence for inbreeding depression and purging of deleterious genetic variation in Indian tigers. Proceedings of the National Academy of Sciences, 118(49), e2023018118.

Kleinman-Ruiz D, Lucena-Perez M, Villanueva B, Fernández J, Saveljev AP, Ratkiewicz M, Schmidt K, Galtier N, García-Dorado A, Godoy JA. 2022. Purging of deleterious burden in the endangered Iberian lynx. Proc. Natl. Acad. Sci. U. S. A. 119:e2110614119.

Kopelman NM, Mayzel J, Jakobsson M, Rosenberg NA, Mayrose I. 2015. Clumpak: a program for identifying clustering modes and packaging population structure inferences across K. Mol. Ecol. Resour. 15:1179–1191.

Korneliussen TS, Albrechtsen A, Nielsen R. 2014. ANGSD: Analysis of Next Generation Sequencing Data. BMC Bioinformatics 15:356.

Korneliussen TS, Moltke I, Albrechtsen A, Nielsen R. 2013. Calculation of Tajima’s D and other neutrality test statistics from low depth next-generation sequencing data. BMC Bioinformatics [Internet] 14. Available from: http://dx.doi.org/10.1186/1471-2105-14-289

Kuang W, Hu J, Wu H, Fen X, Dai Q, Fu Q, Xiao W, Frantz L, Roos C, Nadler T, et al. 2020. Genetic Diversity, Inbreeding Level, and Genetic Load in Endangered Snub-Nosed Monkeys (Rhinopithecus). Front. Genet. 11:615926.

Kumar S, Suleski M, Craig JM, Kasprowicz AE, Sanderford M, Li M, Stecher G, Hedges SB. 2022. TimeTree 5: An Expanded Resource for Species Divergence Times. Mol. Biol. Evol. [Internet] 39. Available from: http://dx.doi.org/10.1093/molbev/msac174

Kuussaari M, Bommarco R, Heikkinen RK, Helm A, Krauss J, Lindborg R, Ockinger E, Pärtel M, Pino J, Rodà F, et al. 2009. Extinction debt: a challenge for biodiversity conservation. Trends Ecol. Evol. 24:564–571.

Labisko J, Bunbury N, Griffiths RA, Groombridge JJ, Chong-Seng L, Bradfield KS, Streicher JW. 2022. Survival of climate warming through niche shifts: Evidence from frogs on tropical islands. Glob. Chang. Biol. 28:1268–1286.

Laikre L, Hoban S, Bruford MW, Segelbacher G, Allendorf FW, Gajardo G, Rodríguez AG, Hedrick PW, Heuertz M, Hohenlohe PA, et al. 2020. Post-2020 goals overlook genetic diversity. Science 367:1083–1085.

Lande R, Shannon S. 1996. THE ROLE OF GENETIC VARIATION IN ADAPTATION AND POPULATION PERSISTENCE IN A CHANGING ENVIRONMENT. Evolution 50:434–437.

Lawson LP, Fessl B, Hernán Vargas F, Farrington HL, Francesca Cunninghame H, Mueller JC, Nemeth E, Christian Sevilla P, Petren K. 2017. Slow motion extinction: inbreeding, introgression, and loss in the critically endangered mangrove finch (Camarhynchus heliobates). Conserv. Genet. 18:159–170.

Li H, Durbin R. 2010. Fast and accurate long-read alignment with Burrows-Wheeler transform. Bioinformatics 26:589–595.

Li H, Durbin R. 2011. Inference of human population history from individual whole-genome sequences. Nature 475:493–496.

Lockwood JL, Cassey P, Blackburn TM. 2009. The more you introduce the more you get: the role of colonization pressure and propagule pressure in invasion ecology. Divers. Distrib. 15:904–910.

Lynch M, Conery J, Burger R. 1995. Mutation Accumulation and the Extinction of Small Populations. Am. Nat. 146:489–518.

Mable BK. 2019. Conservation of adaptive potential and functional diversity: integrating old and new approaches. Conserv. Genet. 20:89–100.

Mallon DP, Jackson RM. 2017. A downlist is not a demotion: Red List status and reality. Oryx 51:605–609.

Mathur S, DeWoody JA. 2021. Genetic load has potential in large populations but is realized in small inbred populations. Evol. Appl. 14:1540–1557.

Mazet O, Rodríguez W, Grusea S, Boitard S, Chikhi L. 2016. On the importance of being structured: instantaneous coalescence rates and human evolution--lessons for ancestral population size inference? Heredity 116:362–371.

Meisner J, Albrechtsen A. 2018. Inferring Population Structure and Admixture Proportions in Low-Depth NGS Data. Genetics 210:719–731.

Monroe MJ, Butchart SHM, Mooers AO, Bokma F. 2019. The dynamics underlying avian extinction trajectories forecast a wave of extinctions. Biol. Lett. 15:20190633.

Nadachowska-Brzyska K, Li C, Smeds L, Zhang G, Ellegren H. 2015. Temporal Dynamics of Avian Populations during Pleistocene Revealed by Whole-Genome Sequences. Curr. Biol. 25:1375–1380.

van Oosterhout C, Speak SA, Birley T, Bortoluzzi C, Percival-Alwyn L, Urban LH, Groombridge JJ, Segelbacher G, Morales HE. 2022. Genomic erosion in the assessment of species extinction risk and recovery potential. bioRxiv [Internet]:2022.09.13.507768. Available from: https://www.biorxiv.org/content/10.1101/2022.09.13.507768v1

Pecnerová P, Garcia-Erill G, Liu X, Nursyifa C, Waples RK, Santander CG, Quinn L, Frandsen P, Meisner J, Stæger FF, et al. 2021. High genetic diversity and low differentiation reflect the ecological versatility of the African leopard. Current Biology [Internet] 31:1862–1871.e5. Available from: http://dx.doi.org/10.1016/j.cub.2021.01.064

Pinto AV, Hansson B, Patramanis I, Morales HE, Oosterhout C. 2022. The evolution of the genetic load during habitat loss and population fragmentation. Available from: https://www.researchsquare.com/article/rs-2123317/latest.pdf

Purcell S, Neale B, Todd-Brown K, Thomas L, Ferreira MAR, Bender D, Maller J, Sklar P, de Bakker PIW, Daly MJ, et al. 2007. PLINK: a tool set for whole-genome association and population-based linkage analyses. Am. J. Hum. Genet. 81:559–575.

Rocha S, Posada D, Harris DJ. 2013. Phylogeography and diversification history of the day-gecko genus Phelsuma in the Seychelles islands. BMC Evol. Biol. 13:3.

Ryan WBF, Carbotte SM, Coplan JO, O’Hara S, Melkonian A, Arko R, Weissel RA, Ferrini V, Goodwillie A, Nitsche F, et al. 2009. Global Multi-Resolution Topography synthesis. Geochemistry, Geophysics, Geosystems [Internet] 10. Available from: http://dx.doi.org/10.1029/2008gc002332

Sánchez-Barreiro F, Gopalakrishnan S, Ramos-Madrigal J, Westbury MV, de Manuel M, Margaryan A, Ciucani MM, Vieira FG, Patramanis Y, Kalthoff DC, et al. 2021. Historical population declines prompted significant genomic erosion in the northern and southern white rhinoceros (Ceratotherium simum). Mol. Ecol. 30:6355–6369.

Santiago E, Novo I, Pardiñas AF, Saura M, Wang J, Caballero A. 2020. Recent Demographic History Inferred by High-Resolution Analysis of Linkage Disequilibrium. Mol. Biol. Evol. 37:3642–3653.

Sax DF, Gaines SD. 2008. Species invasions and extinction: The future of native biodiversity on islands. Proceedings of the National Academy of Sciences 105:11490–11497.

Schubert M, Ermini L, Der Sarkissian C, Jónsson H, Ginolhac A, Schaefer R, Martin MD, Fernández R, Kircher M, McCue M, et al. 2014. Characterization of ancient and modern genomes by SNP detection and phylogenomic and metagenomic analysis using PALEOMIX. Nat. Protoc. 9:1056–1082.

von Seth J, van der Valk T, Lord E, Sigeman H, Olsen R-A, Knapp M, Kardailsky O, Robertson F, Hale M, Houston D, et al. 2022. Genomic trajectories of a near-extinction event in the Chatham Island black robin. BMC Genomics 23:747.

Skotte L, Korneliussen TS, Albrechtsen A. 2013. Estimating individual admixture proportions from next generation sequencing data. Genetics 195:693–702.

Smeds L, Ellegren H. 2022. From high masked to high realized genetic load in inbred Scandinavian wolves. Mol. Ecol. [Internet]. Available from: http://dx.doi.org/10.1111/mec.16802

Smeds L, Qvarnström A, Ellegren H. 2016. Direct estimate of the rate of germline mutation in a bird. Genome Res. 26:1211–1218.

Swaisgood RR, Wang D, Wei F. 2018. Panda Downlisted but not Out of the Woods. Conserv. Lett. 11:e12355.

Taylor SS, Jamieson IG, Wallis GP. 2007. Historic and contemporary levels of genetic variation in two New Zealand passerines with different histories of decline. J. Evol. Biol. 20:2035–2047.

Thompson EA. 2013. Identity by Descent: Variation in Meiosis, Across Genomes, and in Populations. Genetics 194:301–326.

Tilman D, May RM, Lehman CL, Nowak MA. 1994. Habitat destruction and the extinction debt. Nature 371:65–66.

Waples RK, Albrechtsen A, Moltke I. 2019. Allele frequency-free inference of close familial relationships from genotypes or low-depth sequencing data. Molecular Ecology [Internet] 28:35–48. Available from: http://dx.doi.org/10.1111/mec.14954

Warren BH, Strasberg D, Bruggemann JH, Prys-Jones RP, Thébaud C. 2010. Why does the biota of the Madagascar region have such a strong Asiatic flavour? Cladistics 26:526–538.

Watterson GA. 1975. On the number of segregating sites in genetical models without recombination. Theor. Popul. Biol. 7:256–276.

Willi Y, van Buskirk J, Hoffmann AA. 2006. Limits to the Adaptive Potential of Small Populations. Annu. Rev. Ecol. Evol. Syst. 37:433–458.

Willi Y, Kristensen TN, Sgrò CM, Weeks AR, Ørsted M, Hoffmann AA. 2022. Conservation genetics as a management tool: The five best-supported paradigms to assist the management of threatened species. Proc. Natl. Acad. Sci. U. S. A. [Internet] 119. Available from: http://dx.doi.org/10.1073/pnas.2105076119

Willoughby JR, Sundaram M, Wijayawardena BK, Kimble SJA, Ji Y, Fernandez NB, Antonides JD, Lamb MC, Marra NJ, DeWoody JA. 2015. The reduction of genetic diversity in threatened vertebrates and new recommendations regarding IUCN conservation rankings. Biol. Conserv. 191:495–503.

Zhang G, Rahbek C, Graves GR, Lei F, Jarvis ED, Gilbert MTP. 2015. Genomics: Bird sequencing project takes off. Nature 522:34.

