## Supplementary material for "Genetic load and adaptive potential of a recovered avian species that narrowly avoided extinction"

**Table S1 Sample metadata**

| ID | Type | museum ID | Museum | Island | Year | Sex | Total Reads | Coverage | Read Len |
|---|---|---|---|---|---|---|---|---|---|
| SPF1086 | Hist | 1887.12.30.1086 | Cambridge | La Digue | 1880 | M | 137318751 | 4.53 | 48 |
| SPF261 | Hist | 1895.5.1.261 | Cambridge | Seychelles | 1880 | M | 157159682 | 5.17 | 48 |
| SPF262 | Hist | 1895.5.1.262 | Cambridge | Seychelles | 1880 | M | 165408584 | 4.32 | 46 |
| SPF2H | Hist | 1927.12.18.387 | NHM-UK | La Digue | 1888 | M | 173096670 | 6.09 | 53 |
| SPF3H | Hist | 1881.11.14.1 | NHM-UK | Seychelles | 1880 | M | 152921326 | 4.33 | 49 |
| SPF4I | Hist | 1927.12.18.390 | NHM-UK | La Digue | 1888 | F | 114759549 | 3.59 | 51 |
| SPF4J | Hist | 1927.12.18.389 | NHM-UK | Praslin | 1888 | F | 106872254 | 3.4 | 51 |
| SPF4K | Hist | 1927.12.18.386 | NHM-UK | La Digue | 1888 | M | 130099801 | 4.27 | 50 |
| SPF5J | Hist | 1895.5.1.263 | NHM-UK | Seychelles | 1880 | F | 189425205 | 5.94 | 48 |
| SPF5K | Hist | 1927.12.18.388 | NHM-UK | Praslin | 1888 | M | 162549025 | 6.18 | 56 |
| SPF6D | Hist | 1881.11.14.2 | NHM-UK | Seychelles | 1880 | M | 136416796 | 4.9 | 51 |
| SPF6F | Hist | 1881.11.14.3 | NHM-UK | Seychelles | 1880 | F | 161487071 | 5.15 | 51 |
| SPF9 | Hist | 1988.21.9 | NHM-UK | Praslin | 1877 | M | 94230242 | 2.68 | 48 |
| SPF02 | Mod | NA | NA | La Digue | 2007 | M | 37666202 | 9.37 | 171 |
| SPF03 | Mod | NA | NA | La Digue | 2007 | M | 37684407 | 9.4 | 170 |
| SPF04 | Mod | NA | NA | La Digue | 2007 | F | 37647654 | 9.36 | 171 |
| SPF05 | Mod | NA | NA | La Digue | 2007 | M | 37698496 | 9.38 | 171 |
| SPF06 | Mod | NA | NA | La Digue | 2007 | Unk | 37504229 | 9.19 | 172 |
| SPF07 | Mod | NA | NA | La Digue | 2008 | Unk | 38789420 | 9.49 | 171 |
| SPF08 | Mod | NA | NA | La Digue | 2008 | M | 36592547 | 8.97 | 173 |
| SPF09 | Mod | NA | NA | La Digue | 2008 | Unk | 38746275 | 9.46 | 172 |
| SPF10 | Mod | NA | NA | La Digue | 2008 | Unk | 37979707 | 9.36 | 171 |
| SPF11 | Mod | NA | NA | La Digue | 2008 | M | 34750529 | 8.44 | 175 |

| | | | | | | | | | |
|---|---|---|---|---|---|---|---|---|---|
| SPF12 | Mod | NA | NA | La Digue | 2008 | M | 38922148 | 9.55 | 172 |
| SPF13 | Mod | NA | NA | La Digue | 2008 | F | 36390050 | 8.78 | 173 |
| SPF14 | Mod | NA | NA | La Digue | 2008 | Unk | 38550770 | 9.22 | 177 |
| SPF15 | Mod | NA | NA | La Digue | 2008 | Unk | 35139456 | 8.08 | 175 |
| SPF16 | Mod | NA | NA | La Digue | 2008 | F | 39330131 | 9.27 | 176 |
| SPF17 | Mod | NA | NA | La Digue | 2008 | M | 38117012 | 9.01 | 172 |
| SPF18 | Mod | NA | NA | La Digue | 2008 | M | 38711895 | 9.12 | 173 |
| SPF19 | Mod | NA | NA | La Digue | 2008 | M | 37741922 | 8.59 | 177 |
| SPF20 | Mod | NA | NA | La Digue | 2008 | M | 39383969 | 9.72 | 172 |

**Table S2 Diversity loss in bird species.** Loss of genomic diversity of different bird species when compared with historical values from museum-preserved samples. Two different metrics are reported: pairwise nucleotide diversity (Pi) and average heterozygosity (He). Year range and generation range refer to the range of time/generation that spans between historical and contemporary samples. For comparison the values we reported for the Seychelles paradise flycatcher were a 6.4-fold loss in He, 10-fold loss in Pi for a comparison that spans a range of 120-131 years and 60-65 generations (see main text). Methodological differences in how the metrics were calculated (e.g., filtering parameters) could introduce small biases, so while the estimates are not directly comparable, these differences are likely minimal, and the overall patterns should not be compromised

| Species | Generation time | Metric | Delta (fold loss) | Historical sample years | Year range | Generation range | Reference |
|---|---|---|---|---|---|---|---|
| Crested Ibis | 3 | Pi | 1.85 | 1841-1922 | 177-100 years | 56-32 | Feng et al. (2019) |
| Chatham Island black robin | 2 | Pi | 1.8 | 1871-1900 | 143-114 years | 72-57 | von Seth et al. (2022) |
| South Island Saddleback | 8 | He | 4.16 | 1877–1898 | 128-107 years | 16 | Taylor et al. (2007) |
| New Zealand Robin | 4 | He | No significant loss | 1873-1955 | 127-50 years | 31-12 | Taylor et al. (2007) |
| Dutch black grouse | 3 | He | No significant loss | 1893-1953 | 115-55 years | 38-18 | Segelbacher et al. (2014) |
| Mangrove Finch (Isabella) | 1 | He | 1.32 | 1899 | 100-114 years | 100-114 | Lawson et al. (2017) |
| Greater Prarie Chicken | 2 | He | 1.26 | 1951 | 45 years | 22 | Bellinger et al. (2003) |

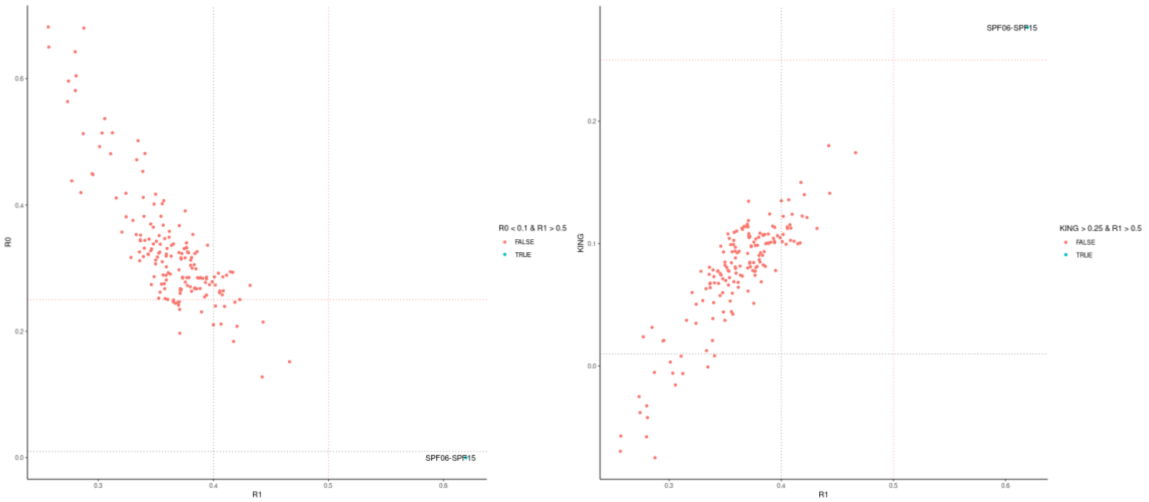


**Figure S1 Genetic relatedness in the modern samples**. We used the method presented by (Waples et al. 2019) which does not rely on allelic frequencies and is suitable for low-coverage data. This method assesses relatedness by comparing two combinations of three kinship statistics making it robust to ascertainment bias: R1 vs R0 and R1 vs KING-robust. We used a low-value threshold for R0 <= 0.1 and high-value thresholds for KING >= 0.25 and R1 >= 0.5 to identify full-sib relationships. We only identified a pair of parent-offspring: SPF06 and SPF15.

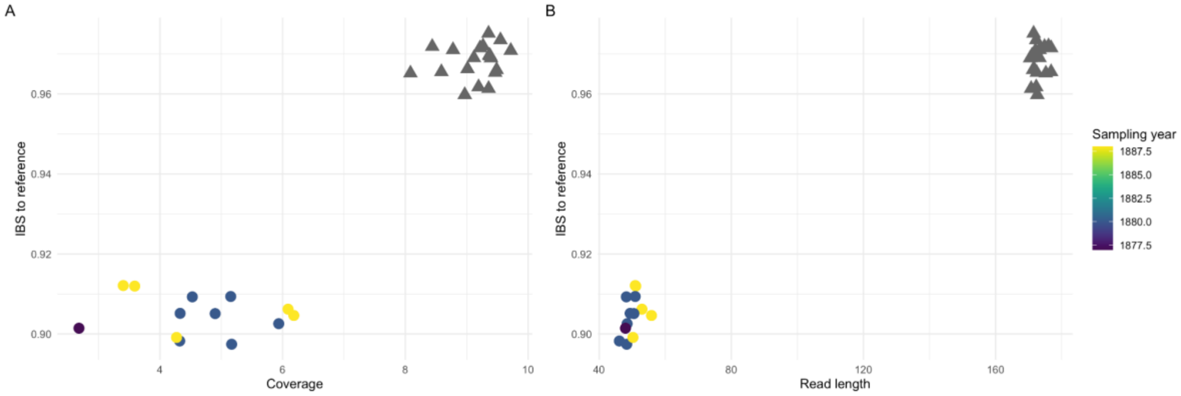


**Figure S2 No evidence of biases introduced by reference genome mapping**. The relatively short average read lengths common to acient DNA (aDNA) caused by post-mortem DNA damage can drive both sequence miss-alignments and a tendency to call SNPs that are biased towards the reference genome (Liu et al. 2021; Gopalakrishnan et al. 2022). Thus, the expectation of mapping reference bias is for the historical samples to have allele calls that are (artificially) more similar to the reference genome, i.e., a smaller identity-by-state (IBS) value. Moreover, low coverage can also create artefacts on the ability to correctly call alleles, with lower coverage disproportionately being called with the reference allele. Here we confirmed that our historical dataset does not suffer from any of these two mapping reference biases as older historical samples (circles) do not have a higher IBS to the reference genome than the modern samples (triangles) and their IBS values are not explained by either (A) coverage or (B) read length. In fact, the relationship is the other way around, with historical samples having a lower IBS than the (modern) reference genome. This is because the modern population has lost a considerable amount of diversity in response to the bottleneck (see main text). To obtain IBS, we randomly sampled one base from the alignments with ANGSD v0.921(Korneliussen et al. 2014) and counted the instances that the allele was the same as the reference genome allele.

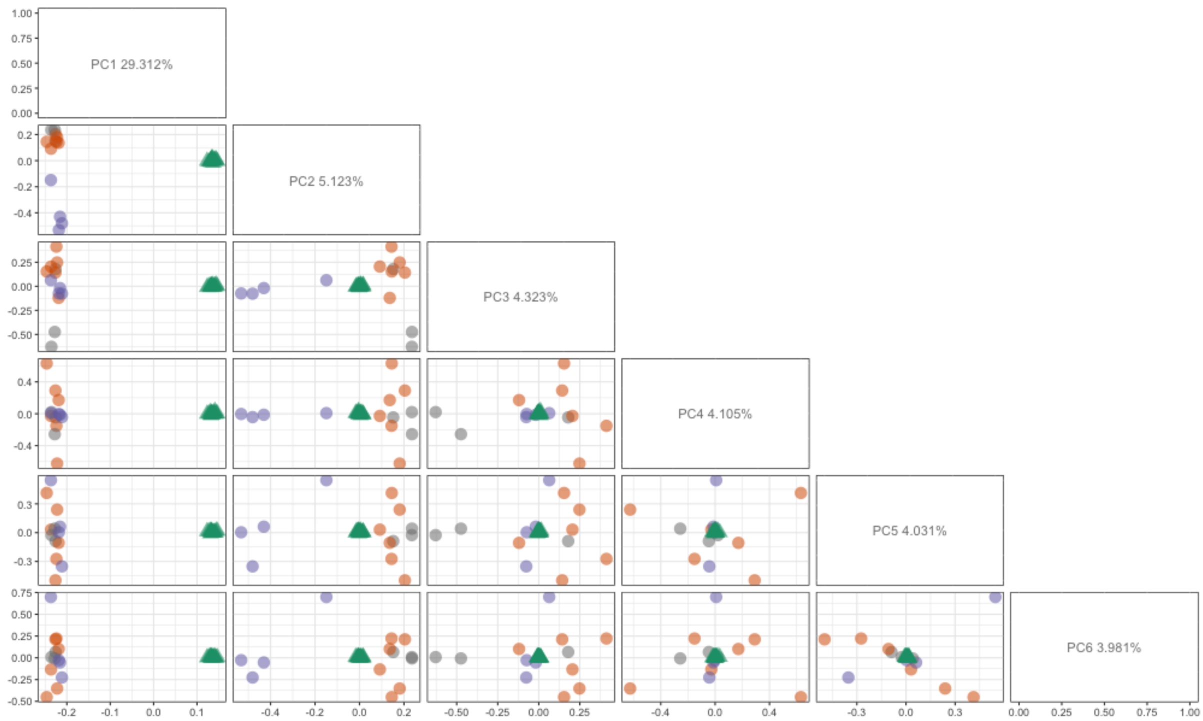


**Figure S3 Principal component analysis of historical (circles; La Digue-orange and Praslin-purple) and modern (triangles; La Digue-green).** One of the challenges of working with old museum-preserved samples is that their associated metadata often lacks precise information about their provenance. In our case, 6 samples lacked information on which island the individuals were collected on. However, using the clear signal of population structure from the PCA we were able to assign those historical individuals to their respective islands.

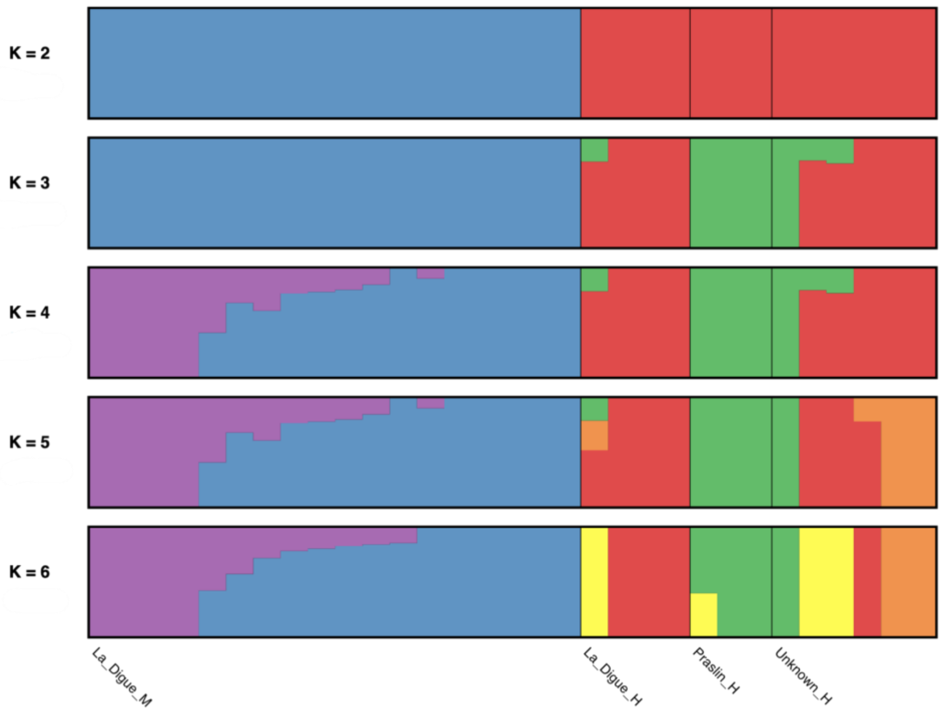


**Figure S4 Admixture analysis.** Visualization of admixture proportions of Modern (La_Digue_M) and Historical (La_Digue_H, Praslin_H, Unknown_H) samples of the best run per K, from K=2-6. Unknown_H groups historical samples with missing sampling location metadata. Colours represent ancestry components. Each bar represents a sample, and samples are sorted by the component with the highest proportion in the group in the highest K. Sample order is the same across all Ks.

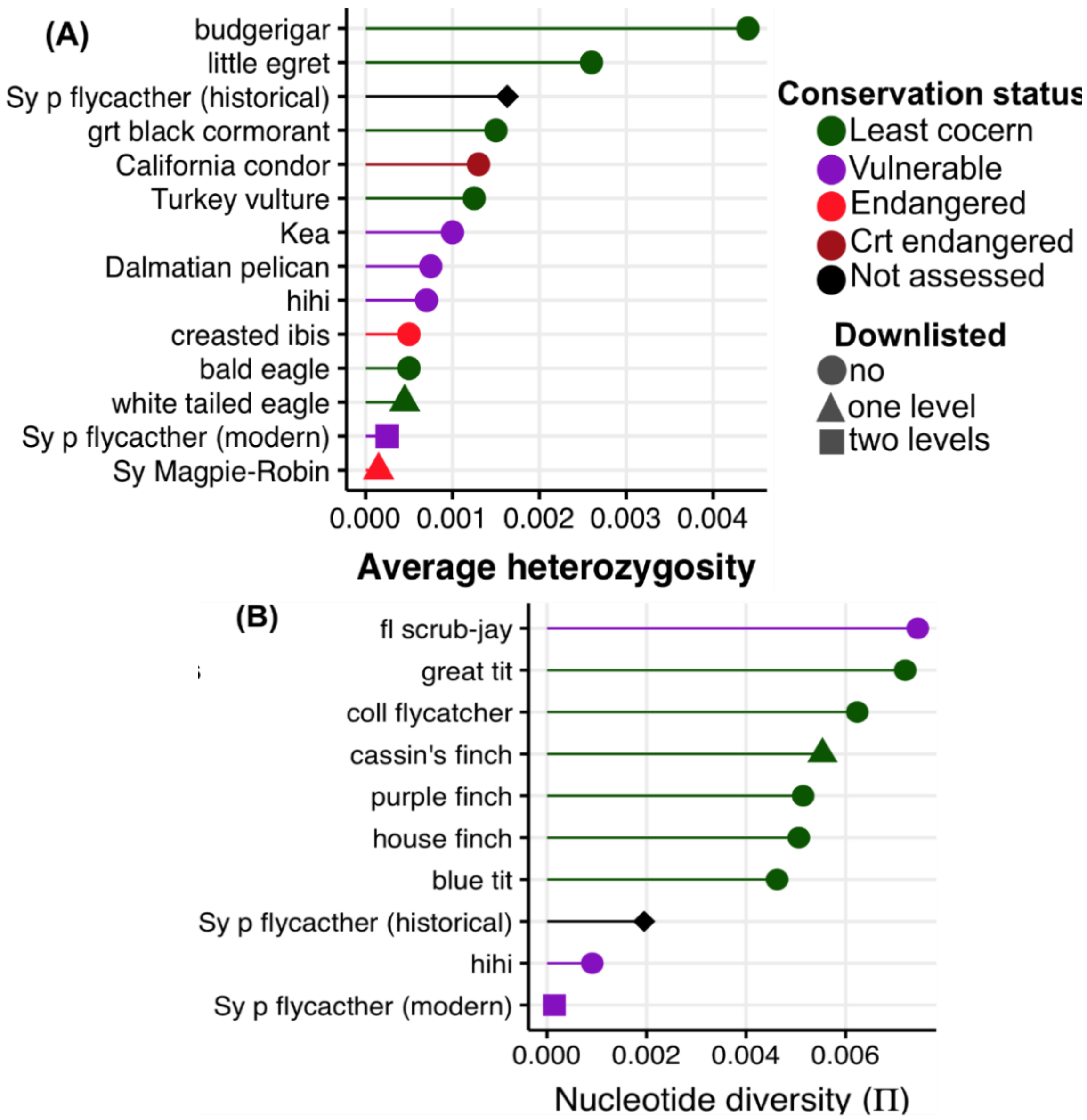


**Figure S5 Genetic diversity for 12 avian species in comparison to the historical and modern estimates for the Seychelles paradise flycatcher** Colours denote conservation status and the symbol denotes how many steps the species has been downlisted in the IUCN Red List. (A) Average genome-wide average heterozygosity. The figure was adapted from (Cavill et al. 2022). Data for nine is from (Li et al. 2014), one from (de Villemereuil et al. 2019) and one from (Robinson et al. 2021). (B) Average genome-wide nucleotide diversity (π). The figure was adapted from (de Villemereuil et al. 2019) with data from (Chen et al. 2014; Shultz et al. 2016; Dutoit et al. 2017; Charles Perrier et al. 2018; C. Perrier et al. 2018). Methodological differences in how the metrics were calculated (e.g., filtering parameters) could introduce small biases, so while the estimates are not directly comparable, these differences are likely minimal, and the overall patterns should not be compromised.

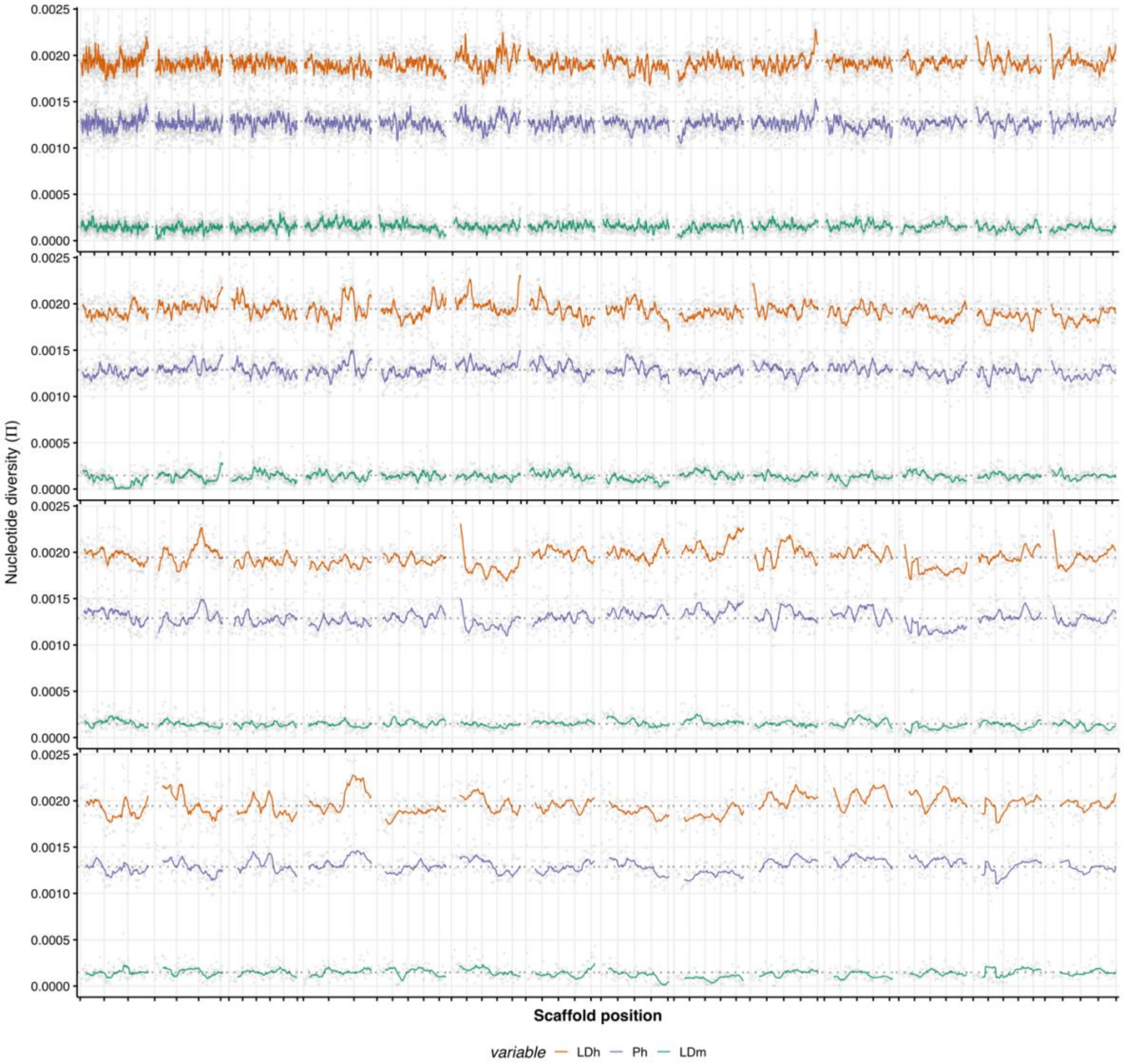


**Figure S6 Nucleotide diversity landscape for historical (La Digue-LDh-orange and Praslin-Ph-purple) and modern (La Digue-LDm-green) populations.** The panels represent continuous runs of scaffolds ordered from long to short. We estimated genome-wide nucleotide diversity (π) from the population-level folded Site Frequency in ANGSD (REF) following Korneliussen et al. (2013) in non-overlapping sliding windows of 50 Kb. Here we plot the 59 autosomal scaffolds longer than 4Mb, which represent ~60% of the total reference genome length

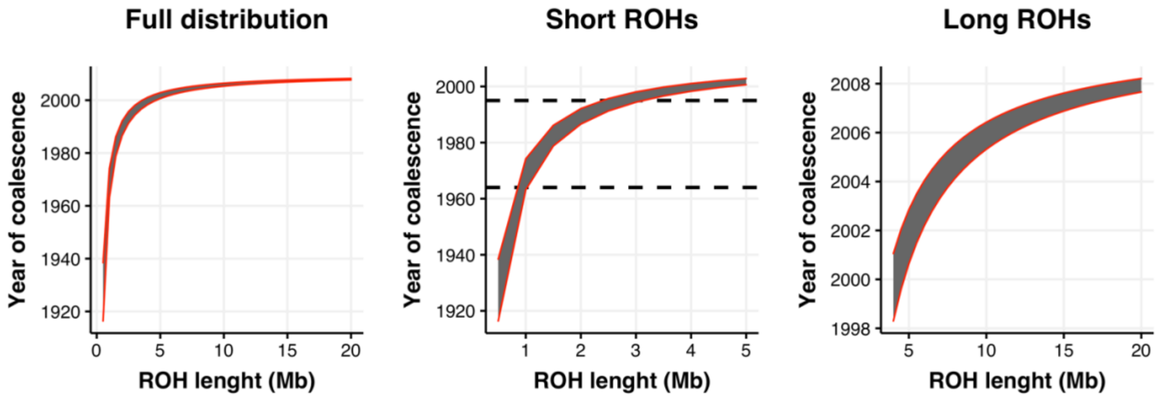


**Figure S7 Expected coalescence time for different Runs of Homozygosity (ROH) lengths.** ROHs are formed when very closely related individuals mate (i.e., inbreeding) and inherit identical segments of DNA to their offspring. As time passes, and if the rate of inbreeding reduces, ROHs are "broken down" by recombination and become shorter. Thus, by estimating the length of a ROH and assuming a constant recombination rate we can calculate their coalescence time, i.e. the time at which the ROH was likely formed. We used the formula *L = 100/2t cM* (Thompson 2013) where *L* is the length of the ROH, *cM* is the recombination rate and *t* is the unknown time of coalescence in generations. We used a range of recombination rate values 1.71- 3.56 cM/Mb from different bird species (collared flycatcher: (Kawakami et al. 2017); zebra finch: (Backström et al. 2010) and Helmeted honeyeater: (Robledo-Ruiz et al. 2022)). We converted generation-ago estimates to years assuming an average generation time of 2 for the Seychelles paradise flycatcher (R. Bristol unpublished). (A) Coalesce times for ROHs length 0.5-20 Mb (B) Coalesce times for short ROH <5 Mb. The approximate time of the bottleneck is indicated in a dashed line. ROHs that formed during the bottleneck should be 1-2 Mb long. (B) Coalesce times for long ROHs >5 Mb that should be formed due to recent inbreeding. In all panels, the shading indicates the 95% confidence intervals obtained using the 1.71- 3.56 cM/Mb range of recombination rates.

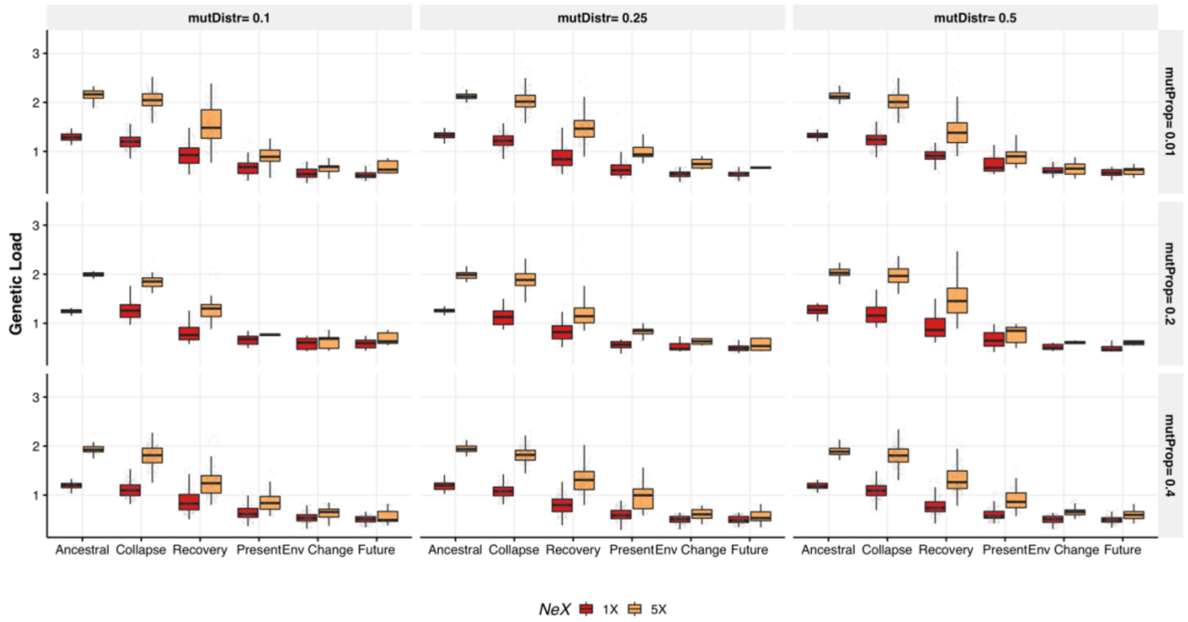


**Figure S8 Parameter test of forward simulations for the (Total) genetic load (masked + realised load).** The 1X trend (red) represents the known trajectory and the alternative scenario represents a 5X (yellow) larger ancestral population size. The panels represent alternative parameters. The parameters *mutDistr* is the range of the uniform distribution from which genotype values (z) were drawn for the polygenic trait (-01-0.1, -0.25-0.25 or -0.5-0.5). The parameter *mutProp* is the relative proportion of mutation contributing to the polygenic trait relative to those contributing to the unconditional genetic load.

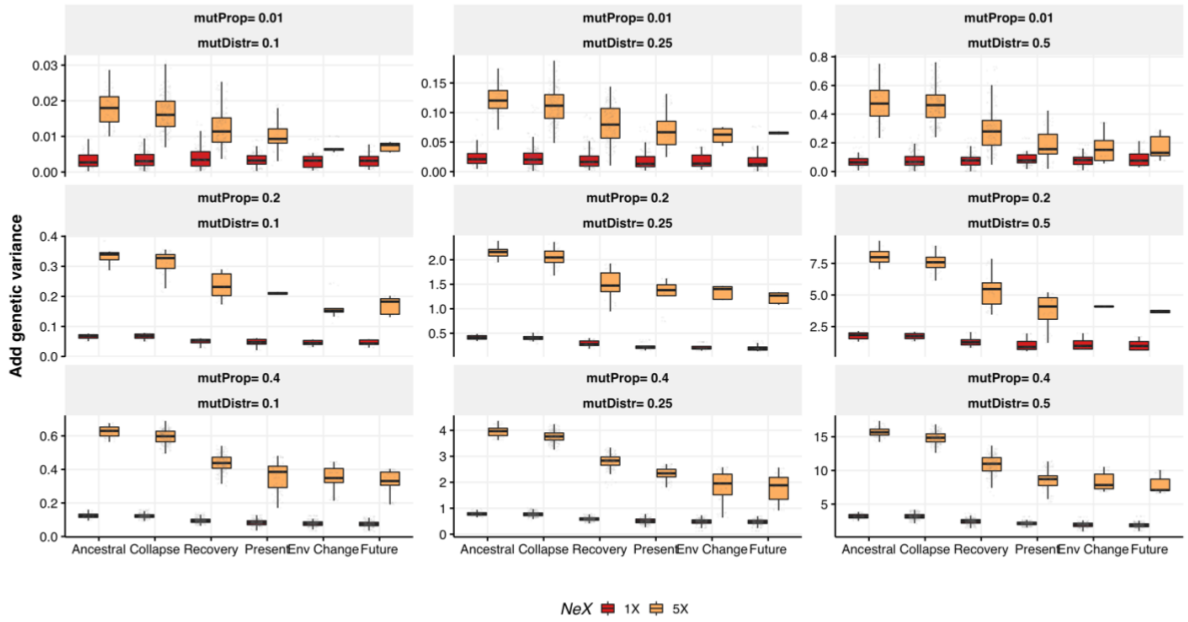


**Figure S9 Parameter test of forward simulations for the additive genetic variance in the quantitative trait (Va).** The 1X trend (red) represents the known trajectory and the alternative scenario represents a 5X (yellow) larger ancestral population size. The panels represent alternative parameters. The parameters *mutDistr* is the range of the uniform distribution from which genotype values (z) were drawn for the polygenic trait (-01-0.1, -0.25-0.25 or -0.5-0.5). The parameter *mutProp* is the relative proportion of mutation contributing to the polygenic trait relative to those contributing to the unconditional genetic load.

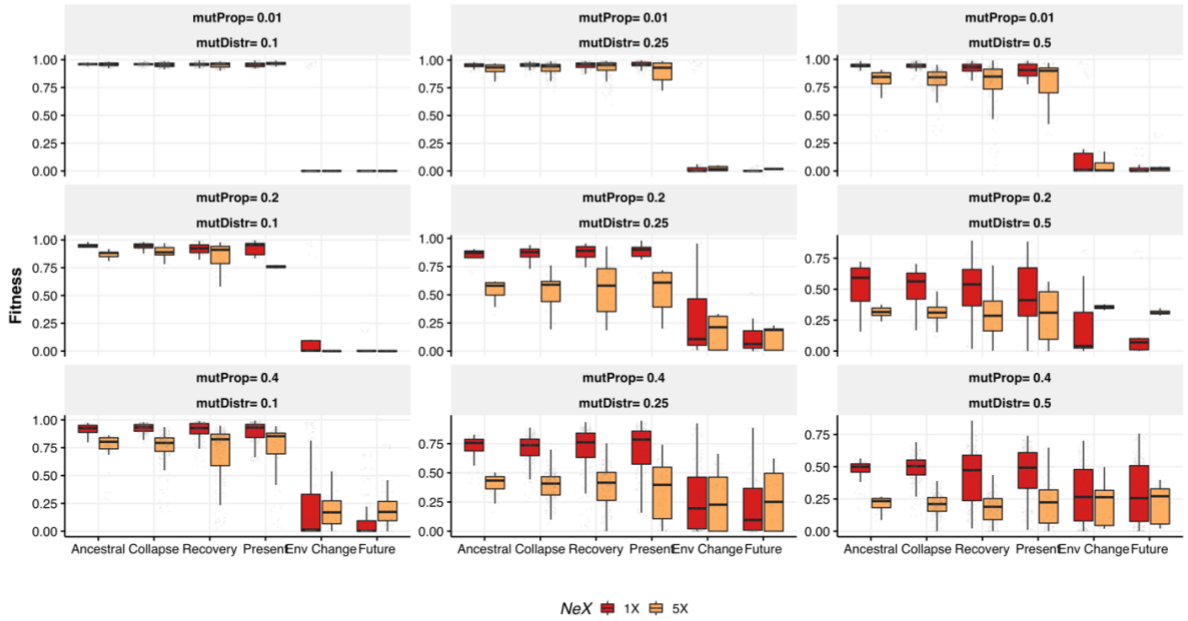


**Figure S10 Parameter test of forward simulations for the fitness effect conferred by the quantitative trait.** The 1X trend (red) represents the known trajectory and the alternative scenario represents a 5X (yellow) larger ancestral population size. The panels represent alternative parameters. The parameters *mutDistr* is the range of the uniform distribution from which genotype values (z) were drawn for the polygenic trait (-01-0.1, -0.25-0.25 or -0.5-0.5). The parameter *mutProp*

is the relative proportion of mutation contributing to the polygenic trait relative to those contributing to the unconditional genetic load.

**References**


Backström N, Forstmeier W, Schielzeth H, Mellenius H, Nam K, Bolund E, Webster MT, Ost T, Schneider M, Kempenaers B, et al. 2010. The recombination landscape of the zebra finch Taeniopygia guttata genome. *Genome Res.* 20:485–495.

Bellinger MR, Johnson JA, Toepfer J, Dunn P. 2003. Loss of genetic variation in greater prairie chickens following a population bottleneck in Wisconsin, U.s.a. *Conserv. Biol.* 17:717–724.

Cavill EL, Gopalakrishnan S, Puetz LC, Ribeiro ÂM, Mak SST, da Fonseca RR, Pacheco G, Dunlop B, Accouche W, Shah N, et al. 2022. Conservation genomics of the endangered Seychelles Magpie-Robin ( *Copsychus sechellarum* ): a unique insight into the history of a precious endemic bird. *Ibis* 164:396–410.

Chen N, Van Hout CV, Gottipati S, Clark AG. 2014. Using Mendelian inheritance to improve high-throughput SNP discovery. *Genetics* 198:847–857.

Dutoit L, Burri R, Nater A, Mugal CF, Ellegren H. 2017. Genomic distribution and estimation of nucleotide diversity in natural populations: perspectives from the collared flycatcher (Ficedula albicollis) genome. *Mol. Ecol. Resour.* 17:586–597.

Feng S, Fang Q, Barnett R, Li C, Han S, Kuhlwilm M, Zhou L, Pan H, Deng Y, Chen G, et al. 2019. The Genomic Footprints of the Fall and Recovery of the Crested Ibis. *Curr. Biol.* 29:340–349.e7.

Gopalakrishnan S, Ebenesersdóttir SS, Lundstrøm IKC, Turner-Walker G, Moore KHS, Luisi P, Margaryan A, Martin MD, Ellegaard MR, Magnússon ÓÞ, et al. 2022. The population genomic legacy of the second plague pandemic. *Curr. Biol.* [Internet]. Available from: http://dx.doi.org/10.1016/j.cub.2022.09.023

Kawakami T, Mugal CF, Suh A, Nater A, Burri R, Smeds L, Ellegren H. 2017. Whole-genome patterns of linkage disequilibrium across flycatcher populations clarify the causes and consequences of fine-scale recombination rate variation in birds. *Mol. Ecol.* 26:4158–4172.

Korneliussen TS, Albrechtsen A, Nielsen R. 2014. ANGSD: Analysis of Next Generation Sequencing Data. *BMC Bioinformatics* 15:356.

Korneliussen TS, Moltke I, Albrechtsen A, Nielsen R. 2013. Calculation of Tajima's D and other neutrality test statistics from low depth next-generation sequencing data. *BMC Bioinformatics* [Internet] 14. Available from: http://dx.doi.org/10.1186/1471-2105-14-289

Lawson LP, Fessl B, Hernán Vargas F, Farrington HL, Francesca Cunninghame H, Mueller JC, Nemeth E, Christian Sevilla P, Petren K. 2017. Slow motion extinction: inbreeding, introgression, and loss in the critically endangered mangrove finch (Camarhynchus heliobates). *Conserv. Genet.* 18:159–170.

Li S, Li B, Cheng C, Xiong Z, Liu Q, Lai J, Carey HV, Zhang Q, Zheng H, Wei S, et al. 2014. Genomic signatures of near-extinction and rebirth of the crested ibis and other endangered bird species. *Genome Biol.* 15:557.

Liu S, Westbury MV, Dussex N, Mitchell KJ, Sinding M-HS, Heintzman PD, Duchêne DA, Kapp JD, von Seth J, Heiniger H, et al. 2021. Ancient and modern genomes unravel the evolutionary history of the rhinoceros family. *Cell* 184:4874–4885.e16.

Perrier C, Delahaie B, Charmantier A. 2018. Heritability estimates from genomewide relatedness matrices in wild populations: Application to a passerine, using a small sample size. *Mol. Ecol. Resour.* 18:838–853.

Perrier C, Lozano del Campo A, Szulkin M, Demeyrier V, Gregoire A, Charmantier A. 2018. Great tits and the city: Distribution of genomic diversity and gene-environment associations along an urbanization gradient. *Evol. Appl.* 11:593–613.

Robinson JA, Bowie RCK, Dudchenko O, Aiden EL, Hendrickson SL, Steiner CC, Ryder OA, Mindell DP, Wall JD. 2021. Genome-wide diversity in the California condor tracks its prehistoric abundance and decline. *Curr. Biol.* 31:2939–2946.e5.

Robledo-Ruiz DA, Gan HM, Kaur P, Dudchenko O, Weisz D, Khan R, Lieberman Aiden E, Osipova E, Hiller M, Morales HE, et al. 2022. Chromosome-length genome assembly and linkage map of a critically endangered Australian bird: the helmeted honeyeater. *Gigascience* [Internet] 11. Available from: http://dx.doi.org/10.1093/gigascience/giac025

Segelbacher G, Strand TM, Quintela M, Axelsson T, Jansman HAH, Koelewijn H-P, Höglund J. 2014. Analyses of historical and current populations of black grouse in Central Europe reveal strong effects of genetic drift and loss of genetic diversity. *Conserv. Genet.* 15:1183–1195.

von Seth J, van der Valk T, Lord E, Sigeman H, Olsen R-A, Knapp M, Kardailsky O, Robertson F, Hale M, Houston D, et al. 2022. Genomic trajectories of a near-extinction event in the Chatham Island black robin. *BMC Genomics* 23:747.

Shultz AJ, Baker AJ, Hill GE, Nolan PM, Edwards SV. 2016. SNPs across time and space: population genomic signatures of founder events and epizootics in the House Finch (Haemorhous mexicanus). *Ecol. Evol.* 6:7475–7489.

Taylor SS, Jamieson IG, Wallis GP. 2007. Historic and contemporary levels of genetic variation in two New Zealand passerines with different histories of decline. *J. Evol. Biol.* 20:2035–2047.

Thompson EA. 2013. Identity by Descent: Variation in Meiosis, Across Genomes, and in Populations. *Genetics* 194:301–326.

de Villemereuil P, Rutschmann A, Lee KD, Ewen JG, Brekke P, Santure AW. 2019. Little Adaptive Potential in a Threatened Passerine Bird. *Curr. Biol.* 29:889–894.e3.

Waples RK, Albrechtsen A, Moltke I. 2019. Allele frequency-free inference of close familial relationships from genotypes or low-depth sequencing data. *Molecular Ecology* [Internet] 28:35–48. Available from: http://dx.doi.org/10.1111/mec.14954
